# Prolonged *in-vivo* tracking of vitreous fluid and its early diagnostic imaging biomarkers for cancer growth

**DOI:** 10.1101/2023.02.15.528694

**Authors:** Mayank Goswami, Snehlata Shakya, Pengfei Zhang

## Abstract

**Purpose:** Estimation of a correlation between cells in vitreous humour and growth in glioblastoma xenografts.

**Methods:** Streams of cells in vitreous humor are observed in optical coherence tomography (OCT) imaging data of animal (NSGS and Athymic Nude-Foxn1^nu^) eyes (34 in total) subjected to xenograft growth study, in-vivo. The cancer disease model is studied with and without nanodrug-based treatment protocols.

**Results:** The presence of CD8+ and CD4+ is reported inside the tumor using the same data earlier. The transition of these cells is shown to take place from the optic nerve via the vitreous into the nerve fiber layer (NFL) at tumor locations and xenograft -related injuries. Functional analysis of dense temporal imaging series (varying from 28 to more than 100 days) reveals a mild correlation between the volumetric growth of the tumor with the density of these cells, quantitatively and qualitatively.

The cross-correlation analysis indicates imaging assisted photodynamic treatment protocol perform relatively better if started with certain delay. Doxorubicin treatment to Nu/Nu Male and NSGS female transforms mild weak negative correlation into mild weak positive correlation.

**Conclusions:** The plots indicate that the mix of the cells in vitreous humor are effectively dominated by immunosuppressor cytotoxic component.

**Translational Relevance:** We propose that the vitreous cell density can be used as imaging biomarker helpful for clinicians in early diagnosis and treatment planning of similar disease models. The limitation of this work is that high resolution OCT systems and data-dependent image segmentation methods are required.

## 1. Introduction

### 1.1. Cells in vitreous humor

The vitreous solutes profile depends upon age, origin (the retina, ciliary body, lens, retinal pigmented epithelium (RPE), or systemic circulation), or ocular pathology^1,2^. Vitreous’ biochemical properties of a healthy eye inhibit cellular migration and proliferation^3^.

In a normal eye, irrespective of genotype of a mouse hyalocytes and microglia are already present inside vitreous cortex just above the internal limiting membrane (ILM) but in relatively negligible density. The innate and adaptive immune response, despite existing immune privilege, may induce or suppress infiltration^4^ of cytotoxic lymphocytes (large granular natural killer (NK) cells ^5,6^, reactive T cells, malignant B cells and associated cytokines arrays), myeloid cells^6,7^, phagocytes (stromal cells, monocytes, histiocytes^8^ and macrophages and associated cytokines), reactive oxygen species(ROS), metabolite, various growth factors, and neutrophils (defensins)^9,10^ and leukocytes^11^. The mutual interaction of these co-existing cells may dysregulate each others’ individual effect around the tumor microenvironment. For example, immunosuppressive cell such as regulatory T cells and myeloid cells are shown to inhibit NK cells’ immunosurveillance and immune effector T-cells’ cytotoxicity ^6^, in-vitro.

Proteomics analysis can categorize more than 1000 different proteins in vitreous, associating certain proteins with a specific ocular disease as a biomarker^2^. Very few in-vivo studies have shown the rare presence of any of these cells in vitreous humor in natural or disease induced conditions^5^.

### 1.2. Etiology

Early diagnosis and etiology (infectious or non-infectious) of acute retinal necrosis (ARN), especially severe cases of posterior uveitis or retinitis and several other ocular diseases, is not easy. It is partly due to ambiguous ophthalmoscopic appearances^12–14^. Diagnosis of ocular reticulum cell sarcoma and myopic choroidal neovascularization, etc., similar to ARN, involves analysis of immunoglobulins from intraocular fluids^15,16^. The pathological reason for a visible increase in vitreous protein concentration, one of the symptoms, can either be inflammation (due to infection, injury, or an autoimmune or inflammatory disease) or from the breakdown of the blood-ocular barrier ^11,17–20^. Vitreous diagnosis helps in studying neoplastic diseases and retinal vasculopathy (for example, diabetic retinopathy)^21^. Cells of noninflammatory origin are studied to investigate uveitis masquerade syndrome (UMS)^22^. Mast cells are also observed in the optic nerve parenchyma with recent severe trauma^23^.

For involved analysis, dense time series of direct images may provide better insight. It may, however, not be practically possible to acquire in the clinical environment. Foremost, it requires non-invasive techniques and a good patient follow-up. Ethically, the priority to implement treatment (for swift patient recovery) suppresses any chance of tracking the natural and very slow-progressing symptoms of ocular diseases.

Laboratory experiments using an apt animal disease model may supplement such desirable analysis with the added advantage of the possibility of obtaining images with higher temporal resolution^24^. Ocular xenografts, or drugs (for example, streptozotocin for diabetic retinopathy) are used for artificial disease induction. Xenograft allows patient-derived contrast-enhanced tissue/cells implantation locally, with relatively better efficiency. However, it is an invasive procedure. Performed once, it can be utilized to supplement in-vivo, non-invasive studies. Xenograft injection (to the retina) for induction or exudation (of vitreous humor) for ex-vivo analysis is a delicate process and requires expertise. It may cause minor puncturing injuries, triggering the inherent biological response mechanism^25^.

### 1.3. Diagnostics

Vitreous protein analysis can be either performed using Ex-vivo and/or In-Vivo diagnostics, both having their own limitations. Ethical issues may pose a constraint against humor sampling for proteomic analysis or saving ocular images to misutilize later for biometrics frauds/identity theft.

#### 1.3.1. Invasive / Ex-Vivo Diagnostics

Although vitreous biomarkers of primary vitreoretinal lymphoma are developed and for uveal melanoma, reports are promising, but diagnosis depends on proper handling of the specimens, methods of aspiration, concentration, fixation, and staining^21^. The immune response of the infiltrating immunoglobulins (clinical and animal models, both) is studied using flow cytometry and immunohistochemistry (IHC) techniques^26^. Exudation of the vitreous humor (from undiluted aqueous humor) requires pars plana vitrectomy/anterior chamber tap/vitreous aspiration tap^12,27^. Polymerase chain reactions (PCR) and Goldmann–Witmer coefficient (GWC) are a few referred diagnostic techniques that assess the intraocular fluid^12,14,18,28^. However, sensitivity to detect specific infections by such tests varies in wide ranges, 46-90% and 25-90% for PCR and GWC, respectively^12^. Flow cytometry analysis, western blotting, and conventional histological studies are other alternatives^11,29,30^. liquid chromatography-tandem mass spectrometry (LC-MS/MS) is used to differentiate the vitreous protein profile^1^. Several other tests are also described here^2^. LASER scanning cytometry and liquid-based cytology also have found utility in cell imaging and analysis.

Sometimes multiple tests are desirable as several patient characteristics (such as patient’s age) affect the choice. More than one such test may not be possible same time due to the relatively lower aqueous humor volume. Multiple extractions via invasive procedures may involve complications^25^. The exudation is a complex microsurgical procedure, especially of the vitreous cortex. It may be time-consuming and fraught with ethical issues. The ex-vivo analysis thus most of the time only facilitates a one-time analysis option. Several clinical studies suffer from poor temporal resolution as the multiple procedures are discouraged. Moreover, invasive tests affect the natural microenvironment and cell dynamics producing inaccurate conclusions. Localizing cells in the vitreous body using an invasive method is reported Accuracy in less as the organ shrivels during the fixation^4^.

#### 1.3.2. In-vivo Non-Invasive Diagnostics

Optical imaging-based non-invasive techniques, namely: LASER Speckle Imaging (LSI) and Fluorescence Correlation Spectroscopy (FCS), are used to explore the dynamic features of subcellular components, cellular motility, cell infiltrations/migrations during inflammation^11,31,32^.

A combined approach using Raman and Holographic microscopy analysis is reported to identify and discriminate the cell populations of CD4+ T cells, monocytes, and B cells^33^.

Fundus and scanning LASER ophthalmoscope (SLO) imaging (Fig. 1(a)) is abundantly used as an affordable alternative and non-invasive tool in both clinical and laboratory applications. It, however, requires contrast enhancement tags (Fig. 1(b)) or may depend on autofluorescence. It gives 2D enface images with image quality significantly depending on optical vitreous transparency. A clinical study is reported observing the transition of cells and other sub-cellular components in-between retinal layers using microscopy ^34^.

**Figure 1.**
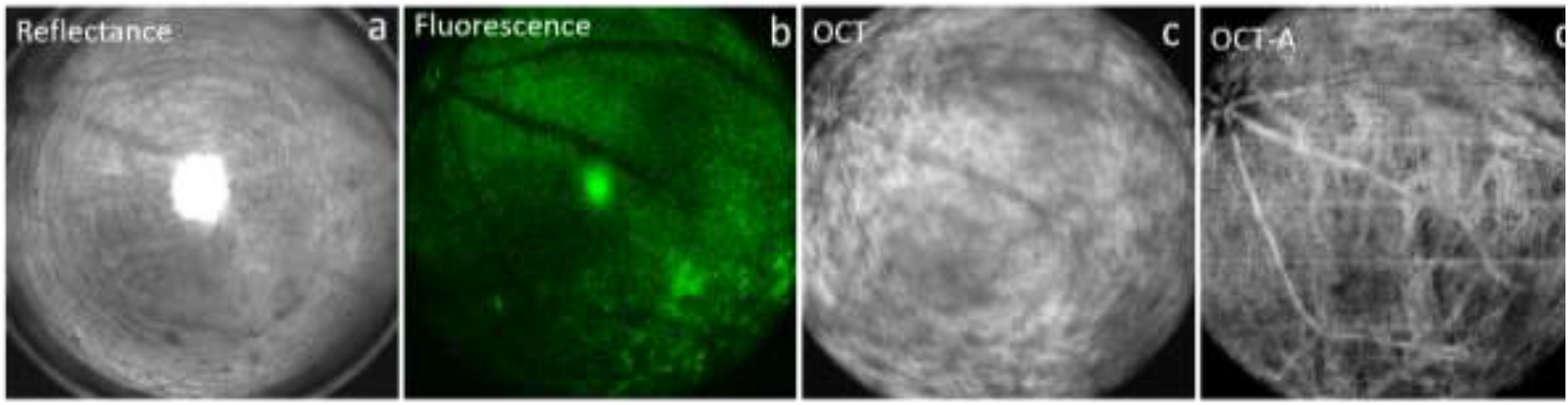
Enface Images of a mouse from a combined OCT animal imaging system. Reflectance and fluorescence are direct fundus images without and with filters. OCT and OCT-A are corresponding images obtained at the same instance.

Optical coherence tomography (OCT) imaging (Fig. 1(c)) is used for similar reporting with relatively better spatial resolution and acquiring speed^16,35^.

Multiple features of OCT have also been used for investigating inflammatory cells for cell sorting in the aqueous chamber as an ex-vivo tool(36,37) and identifying and quantifying macrophages in coronary artery plaque and *in-vivo* immune cell migration^36,37^. Studies also reported that OCT could be used to visualize immune cells and evaluate cellular characteristics and dynamics through rodent models^35,38^. Similar cellular structures are reported in humans^39^.

Optical coherence tomographic-angiography (OCT-A) or phase variance OCT (PV-OCT) provides blood vascular 3D maps with a micron-order spatial resolution^24,36^. The combined/multi-spectrum (SLO-OCT-OCT-A) animal images (shown in Figs. 1(a-d)) provides effective co-localization of 3D ocular structures with 2D fundus enface images. OCT images offer a good match with conventional histological studies^40^.

A study about the artificial retinal detachment(RD) model has reported that monocytes (extravasated from the vasculature and vitreous surface of the retina) as primary immune cell-mediated cytokine storm^41^ causing Interleukin-6 (IL6) signaling in vitreous humor while microglia remained un-effected. One study showed the presence of microglia and hyalocytes distribution using OCT imaging^42^. An in-vivo study (supported by immunohistochemistry and flow cytometry analysis) has revealed infiltration of CD11b^+^ CD45^+^ myeloid cells during photoreceptor cell death (induction model in mouse) across retinal vessels within 48 hours of photoreceptor degeneration^7^.

### 1.4. Other frontiers

Besides needing to record the dense time-dependent cellular activities, several necessary advancements are under development in the field of functional image processing for disease-specific feature segmentation. Several semi-automatic and automatic soft tools are reported ^43–47^. Methods to segment retinal layers are well established except for describing the estimation of average Cell counts in vitreous humor in the vitreous region.

### 1.5. Motivation

The previously published data shows the treatment efficiency of nano doxorubicin on the murine tumor model ^24^. The histopathological analysis shows the existence of CD8+ (an antiviral cytotoxic T-Cell) reported inside the retina alongside the growth of glioblastoma xenograft. The consistent presence of cellular bodies in the vitreous fluid is observed in respective OCT data. The cell motility and density varied with the tumor growth relative to the healthy retina. We present an exploration of the dynamics of these particles quantified along with the tumor progression using the same data set. Primarily, we would want investigation here that if the existence of a cellular body in vitreous fluid: (a) is entirely due to injury caused by the xenograft process and disappears while the retina heals, (b) supports the healing process of the retina, and (c) can be correlated with the growth of the tumor. We also like to explore if the cellular bodies are spewed from the site of injury or tumor or entering from the vitreous fluid into the retina. Functional OCT imaging data is used to estimate the average Cell count in vitreous humor as the main characterization parameter.

The following sections explain methods (in brief) to carry out this study, followed by qualitative analysis, results, and involved discussion.

## 2. Materials and Methods

### 2.1. Small animal husbandry

Mice of two genotypes (a) athymic nude-foxn1^nu^ or Nu/Nu (12 male and 5 female) and (b) immune suppressed NOD.Cg-Prkd^scid^Il2rg^tm1Wjl^/Tg or NSGS (4 female)of 6-8 weeks of ages kept on a 12:12 light cycle are used^48,49^. Tumor growth data is borrowed from earlier published work^24,50^. Nu/Nu are implanted with glioblastoma cells and NSGS are implanted with Patient derived xenograft (PDX) cells.

### 2.2. Xenograft preparation and transplantation

All the cell culture protocols and the methodology of xenograft glioblastoma tumor cell (U87MG-GFP) preparation for transplantation are already reported^24^. Ocular transplantation of xenografts between the retinal pigment epithelium (RPE) and the retinal cells was carried out using the method of Matsumoto et al. ^51^. Mice with retinal holes or sub-retinal or vitreous hemorrhages were excluded from the study.

### 2.3. Imaging-assisted Nano drug treatment

Total of 34 animal eyes are followed. Each animal (starting of experiment when aged between 5-6 weeks old) was followed from 5^th^ day before (as baseline) till the end of the treatment protocol. The Nano drug samples consisting of porphyrin-PEG-doxorubicin termed as ‘nanodox’ were prepared according to Li et al. and administered as reported^52^. Prior to imaging, all the necessary animal handling procedures are taken to keep the mice ready. The optical features and parametric settings of the multimodal imaging systems, Optical coherence tomography (OCT), and scanning laser ophthalmoscopy (SLO) are described in detail^53^.

### 2.4. Immunohistochemistry and quantitative data

Histological studies still remain the efficient biological assay for the confirmation and extended *in vivo* investigation of the tumor. All the protocols followed for conducting histopathology and flow analysis are similarly reported. A few of the available histological results of the processed data are used in the discussion section. Tissue transformation due to immune cells infiltration into the tumor through flow cytometry analysis is also studied^24^.

### 2.5. Cell identification by the deep learning algorithm

The flowchart is shown in Figure 2. Each OCT Volume data contains 359 B-Scan. Entire data set contains total 537064 (359 x 34 (eyes) x 44 (imaging days on-average per eye)) B-Scans. Each volume of data went under the pre-processing stage for registering the data on the same level, contrast stretching (to reduce Tyndall scattering), and resizing / rescaling^17^. Afterward, 10K images were manually annotated for the presence of cells (in vitreous fluid) and NFL. Cells above the NFL are easily visible relative to those cells which migrated inside the retinal layers.

**Figure 2:**
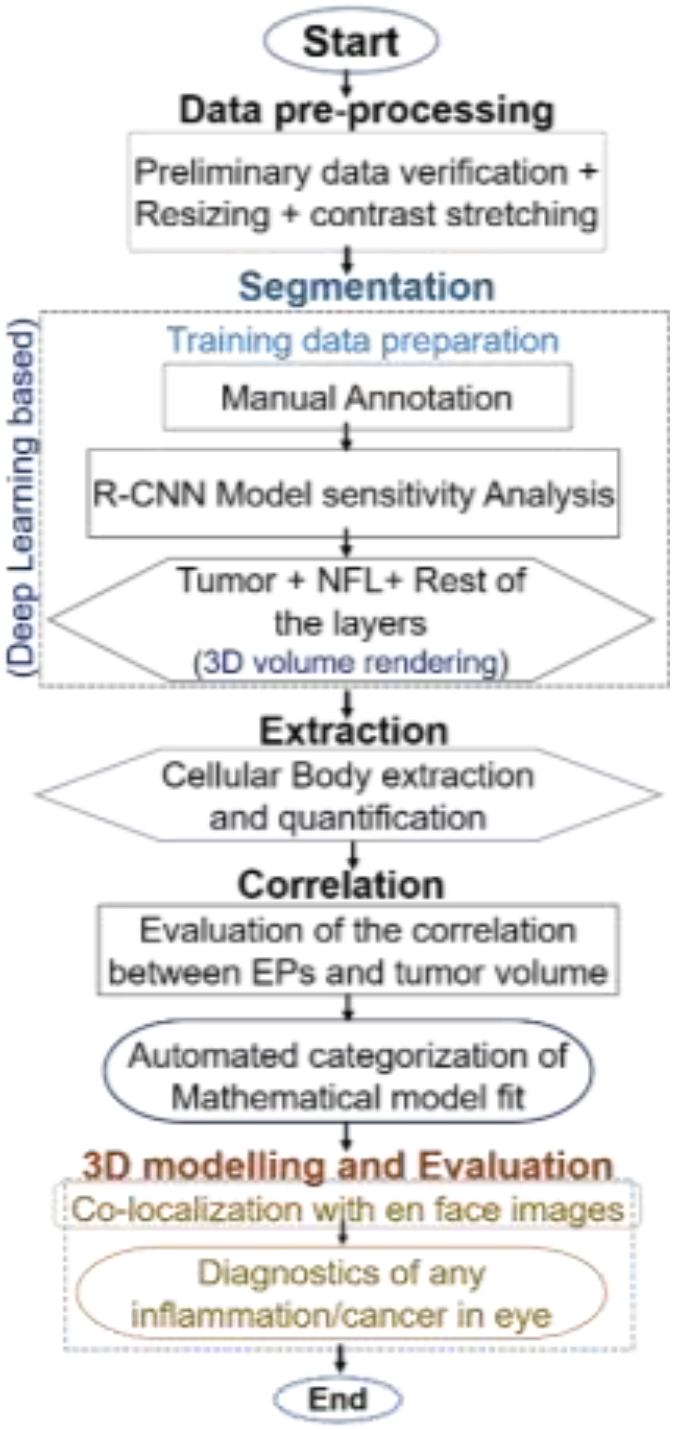
Flow diagram showing the methodology of Image processing and data analysis

The annotated data is used to train a deep learning-based hybrid model (U-net+ResNET50)^46^. The trained model with acceptable performance (Dice Coeff. equals 0.893) is used for locating the cells in all eyes. A single cell having a size larger than the Z resolution of a single B-Scan is present in more than one B-Scan slice. The counting algorithm may count it more than once. The failure to merge all pieces of a single cell from slices can be removed if counting is also performed by an AI algorithm using manually trained data. However, for experts, identifying the number of cells during the annotation, if they are clumped, is impossible. It can only be done by dividing the volume of the clump (of more than one cell: a cell molecule) by the volume of a single cell. In another approach, cells are encapsulated in respective voxels (size 10-20 microns)^54^. It, however, has shown minor differences as compared to the previous method. The overestimation (if any) would be present throughout the data; thus, it is assumed that it will not affect the estimation.

Classical linear regression (with the best fit) estimates the possible correlation between cell density with normalized volumetric tumor growth w.r.t gender and genotype. The extracted and segmented components were augmented using the generalized MATLAB™ code for 3D visualization. Co-localization and validation of 2D B-scan and 3D images with the histology data are done. Finally, the results were analyzed and discussed.

## 3. Results

### 3.1. Immunoglobulin cells, retinal tissues, or collagen fibrils

Standard baseline imaging revealed presence of cells (having relatively faint brightness to other structure) before xenograft. Figures S1 and S2 shows few BScans (on several retinal axial/transverse planes/cross-section) are few of the example. The close observation indicates the proximity of cells with blood vessels in ILM. The dynamics shown in <u>Movie M0</u> inspires to think that stream of cells is originating from these blood vessels and communicating to each other via vitreous humor. These thick blood vessels in ILM are also connected to optical cord.

Micron resolution (2 microns) ocular OCT imaging data (in 3D) may help visualize the viscous humor structures above the retinal surface map, as shown in Figure 3. The discrete nature of these structures indicates dissimilarities with collagen fibrils, vitreous detachment, or hyaloid canal ^55,56^. These structures appear to be more like air pockets^57^. Oval shape surface of red blood cells (RBC), and other similar molecules, migrating between retinal layers or moving inside blood vessels, can be perceived in corresponding OCT-A data (shown in Fig. 3h). It also shows each molecule contains a sheathing layer (shown in a fake white color scheme). We believe it is due to the presence of intermolecular plasma fluid^58,59^.

**Figure 3:**
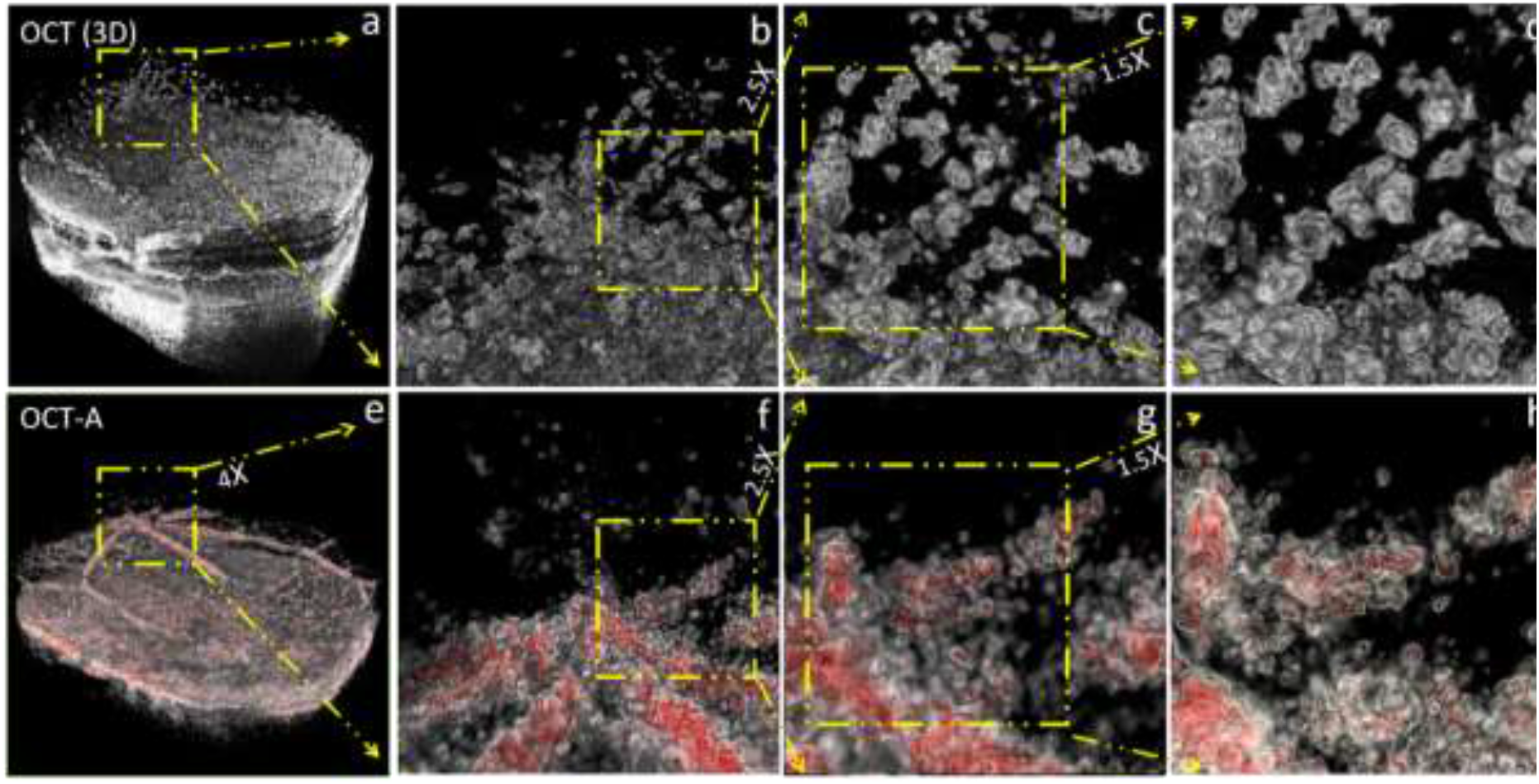
Micro resolution 3D OCT and OCT-A images. After xenograft, the OCT shows a significant presence of cells floating in vitreous fluid, and OCT-A shows an oval shape of red blood cells. (a, e) Retina Volume OCT and OCT-A, (b-d) respective zoom section showing the discrete structures at the optic nerve and surrounding in OCT, (f-g) shows blood vessels using fake red and white color scheme, (h) oval shape red blood cells shown covered what appears to be with white plasma sheath.

The 3D surface map, visible in OCT but not in OCT-A, may help define immunoglobulin cells (in vitreous fluid). If OCT-A images do not show the presence of molecules whereas OCT shows, these molecules are considered immunoglobulin cells (only above the NFL). Few cases can be observed by visually comparing respective figures (row-wise Figs. 3a-3d and Figs. 3e-3h). A supplementary figure S3 shows a composite figure (Fig. 3c merged/overlapped with Fig. 3g), highlighting the existence and absence of structures seen in OCT and OCT-A images.

### 3.2. Qualitative evaluation of cells migration after xenograft

We note that the field of view (53°) of the imaging system is able to capture a limited section of the retina. The data shown in Figures 1, 3, and 4 is of an athymic mouse taken after the second day of xenograft. The site of injury made by a needle is clearly visible (Figs. 4a) with a rhegmatogenous detachment (right top part of the sclera). Right before the xenograft, it was not present; thus, its origin is more likely due to traction created in vitreous humor by the needle’s exertion against its adherence to the retina^57^. Figure 4c shows a few stacks (3D) belonging to the site of injury and distribution of immunoglobulin cells on the top of the retina. The same distribution is shown with and without retina in Figs. 4d and 4e. Figure 4d shows the segmented retina (upper layer NFL till the end of sclera) and all the molecules above it. The B-Scan stacks from the front site of the injury till the end of the retina are shown in Figs. 4f - 4i to highlight the 2D distribution of these cells. The pathway/trajectory (highlighted in Figs. 4a - 4c) of the stream transporting these molecules can also provide a clue about their origin. It is described earlier that these cells may have amoeboid movement^4^.

**Figure 4:**
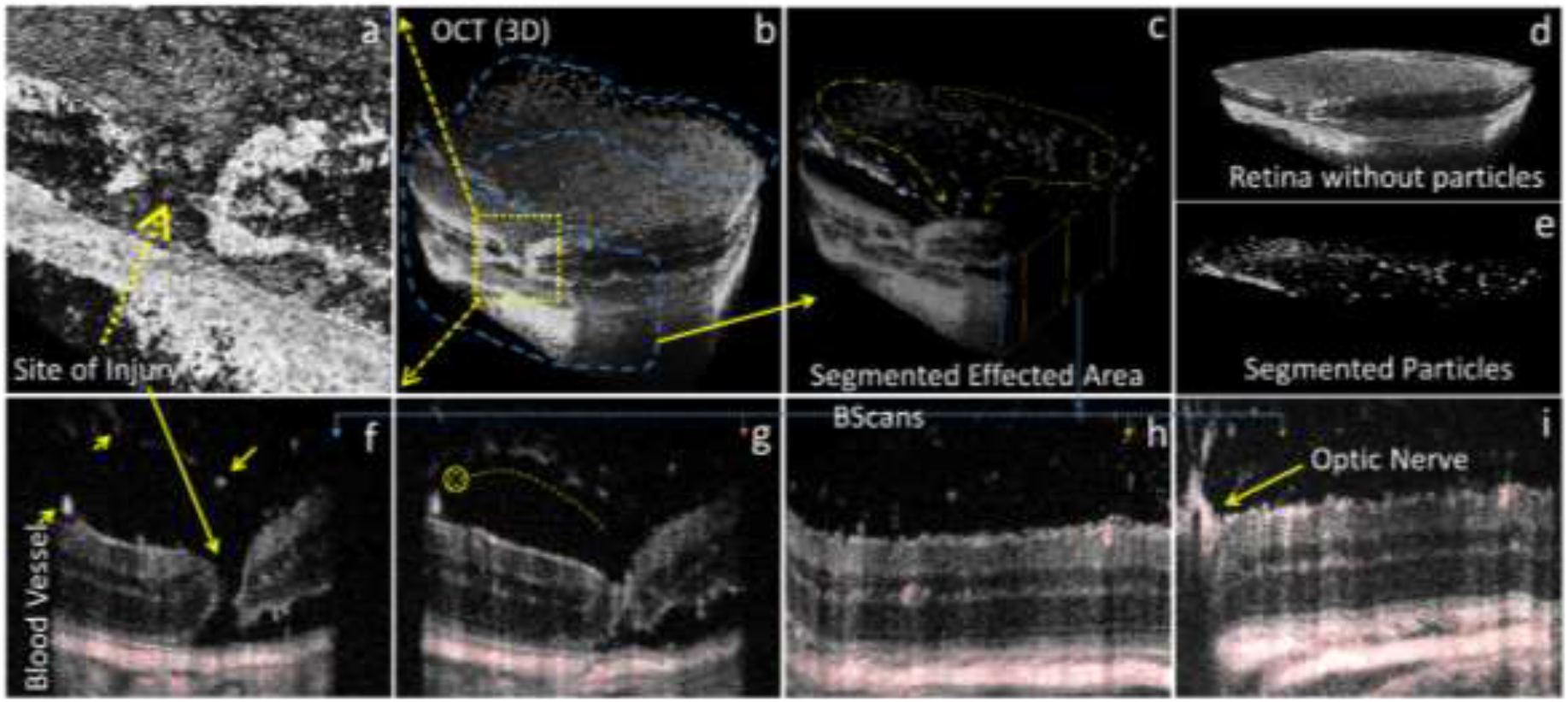
Segmentation of OCT data to estimate immunoglobulin cells density in vitreous fluid; (a) showing the site of injury, (b) full 3D volume, (c) the segmented affected area showing two channels/streams of particles, dense at optic nerve end, one towards the site of injury another towards the detached retina, (d) separate sections of the retina and segmented particles, (f-i) B Scan showing site of injury along with detachment, detachment only, a healthy retina and section cut containing optic nerve. (f) shows particles in fading and bright spots showing fading are in another Y plane traveling inside. The red tint in Fig. 4 (f-i) indicates OCT-A overlapped with OCT.

These floating molecules can either be the dislodged tissues (from the upper layer of the retina) being spewed from or immunoglobulin cells either leaking from or entering the injury and detachment spots (Fig. 4a). The dislodged retinal tissues (from the retinal surface) may have at least few blood cells attached thus expected to remain visible in OCT-A images as well. However, the density of particles in streams of molecules is relatively higher in OCT images as compared to the corresponding OCT-A images (<u>Movie M1</u>) and Figure3. It indicates that by the time of imaging, the particles in the vitreous have lost their driving mechanisms to move, especially red blood cells. On successive days, these OCT-A images show a similar absence on corresponding spots.

There can only be one possibility for these streams to transport the retinal tissues, spewing inside out. Also, the stream must remain till the injury is not healed. However, these cells (lower in density) are found well before xenograft. Figure S3 and <u>Movie M2</u> (supplementary file) show that stream of cells is connected between the optical cord and other blood vessels near to ILM inside the vitreous cortex only. This data is taken before xenografting the tumor cells in female Nu/Nu mice. Table 1 provides the details.

**Table 1.** NSGS

| Condition | Cases with Treatment |  |  | Cases without Treatment |  |  |
| --- | --- | --- | --- | --- | --- | --- |
|  | M | F (4) |  | M | F (4) |  |
|  |  | VH. Cells<br>(x10<sup>6</sup>) | Tumor Vol. |  | VH. Cells<br>(x10<sup>6</sup>) | Tumor Vol. |
| Before Xenograft | x | 3.735 | 0 | x | 3.113 | 0 |
| After Xenograft | x | 4.683 | 3.142 | x | 3.599 | 10.472 |
| Before 1<sup>st</sup> Treat | x | 4.466 | 10.038 | x | x | x |
| Before 2<sup>nd</sup> Treat | x | 3.768 | 10.188 | x | x | x |
| Before Euthanizing | x | 3.391 | 5.343 | x | 3.563 | 13.971 |

\*VH. Cells: Average Cell counts in the vitreous humor, x: data was not available

The stream of molecules (clumped cells) is seen in OCT images right after the xenograft session (images taken after 5^th^ day), strangely connecting the site of injury to the untouched optic nerve head, which is far away from the location of the needle. The bright molecules are in the very plane of B-Scan and what appears to be fading (as we move from the site of injury towards the optic nerve) are in deep Z planes (Figs. 4f-4i). The corresponding 3D distribution is shown in Figs. 3b and 4b. The spread of molecules (between the optic nerve and the site of injury) is into two distinct channels. The centre part near to the optic nerve appears to be in a crown-like shape. We also note, in this data, right after the xenograft, the retina shows slight detachment (right top part of the sclera). The second channel is seen to be associated with it or has risen in symmetry due to vitreous gel dynamics. In other datasets, this second channel is not visible; thus, the later argument seems less plausible.

The density of the floating molecules is higher at the optic nerve side (figs. 3a-3d). The spread indicates that these molecules are segregating away from the optic nerve. It may be possible that tissues originating from the site of injury and detachment are driven towards the optics nerve due to vitreous fluid dynamics as the head of the cord is protruding out^60^. The viscosity of humor is relatively high at the edges than at the center.

We note that the path to clear/flush out the fluid, however, is not towards the optic nerve. Physiologically the major ocular blood supply is via the choroid towards the optic nerve. Alternatively, these molecules are in the bloodstream, flowing via choroid towards the optic nerve and then targeting the injured site by traveling all the way from the optical nerve opening to the site of injury. The latter argument seems least likely as the optic may not have any natural opening to release the molecules into vitreous fluid and is not injured during xenograft either.

The presence of these cells in relatively healthy retina / before xenograft also supports the claim that these are predominantly the immunoglobulin cells being released for the site of injuries from the optics nerve, mainly. It is reported earlier that the optic nerve is an active infiltration site of the inflammatory cell^61–63^.

### 3.3. Tumor Growth and cell density

Once the injury (if present due to xenograft) heals and the tumor proliferates, some of these bright particle-like structures / cells in vitreous humor are seen (in time series of direct OCT images) moving in streams while their distance gets shorten with respect to ILM. Figure 5 shows one particular example taken from a Nu/Nu male mouse that lived more than 100 days. Fig 5(a) shows a small tumor volume (13.5 / 359 (total B-Scans) cancer cells/volume (*μl*)) and a significant accumulation of cells thick yellow arrows) in vitreous fluid. As time progressed, the density of cells in vitreous fluid varied, and so as the size of the tumor (167 / 359 (total B-Scans) cancer cells/volume (*μl*)) in this B-Scan. The density increased further, and multiple streams are observed in Fig. 5(c). This particular retina is subjected to nanodox treatment after 100 days, which affects the tumor volume. Initially, the cancer cell number and tumor volume decreased^24^. At the end of the treatment sessions, cancer cell density is found to be negligible (verified by histopathology results shown in the last section of this paper). Figure 5(d)) shows that after treatment, cell density relatively decreased. Figure 5(e) shows a bloated retinal surface with a large population of vitreous cells. In one of the reported works, the transformation of this tumor into a cyst is proposed^46^. One minor observation following this time series is that most of the time distribution of these cells remains in the vicinity of two blood vessels. These two blood vessels (marked by thin red arrows) are also taken as a point of reference to locate the same B-Scan every time. Supplementary figure S3 is provided with 3D images carrying in-depth visualization. The 3D cell dynamics is described and depicted in fig. S4.

**Figure 5:**
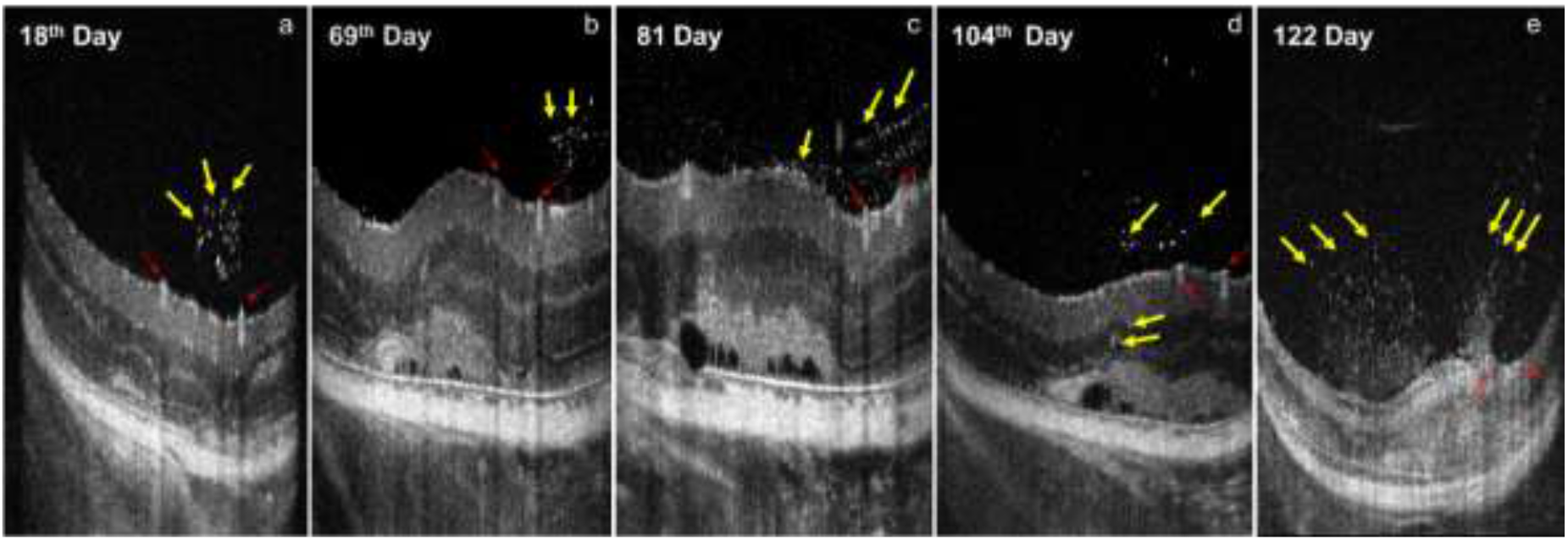
Successive tracking (of same retinal axial /cross-sectional location) of cells in vitreous humor vs. tumor growth shown in the B-scan timeline; (a) cells at the site of inflammation after 18 days of xenograft with small tumor at this B-Scan Slice, (b) relatively few immune cells after 69^th^ days as the tumor got bigger, (c) tumor detachment at multiple sites after significant growth with multiple cell streams, (d) some of the cells entering inside the retina towards tumor and (e) immune cells infiltration into tumor after 122 days. Yellow thick arrows are markers for cells, and thin arrows (in red) are the point at blood vessels as landmarks showing all B-Scans are taken approximately on the same location.

### 3.4. Immune cells: A natural prognostic biomarker for early diagnosis

OCT images including all possible cases: (a) eye in healthy condition with and without the presence of the cellular body, (b) before and immediately after xenograft injury, (c) several days after xenograft with and without nano-dox treatment are analysed, and presented now. The eyes having Cells in vitreous humor below this threshold value (when Tumor Volume is zero, i.e., before xenograft) are considered healthy. The data in Tables 1 and 2 contains average values of all cases, i.e., with and without treatment. Figures S5 and S6 contain full data. The study lacks NSGS Male mice data.

**Table 2.**
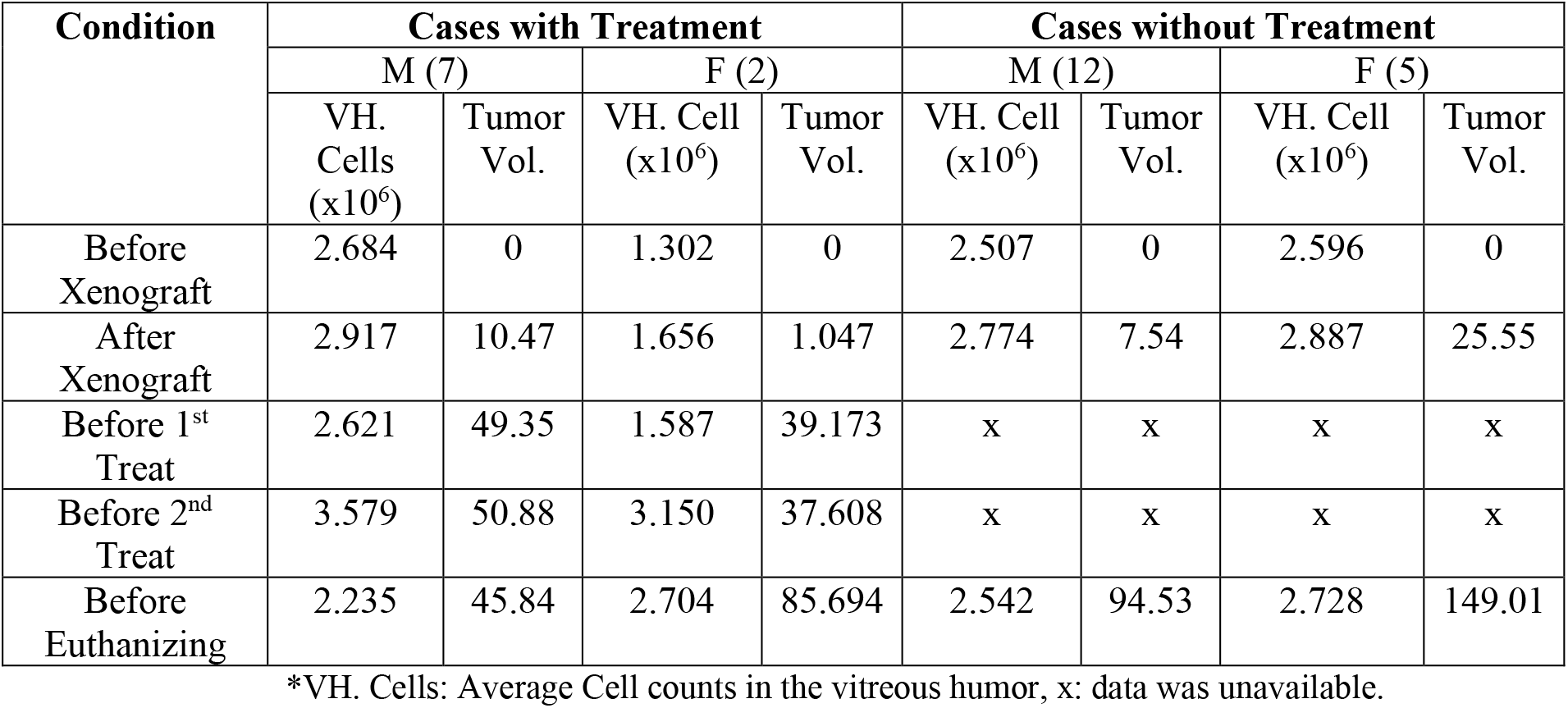
Nu/Nu

| Condition | Cases with Treatment |  |  |  | Cases without Treatment |  |  |  |
| --- | --- | --- | --- | --- | --- | --- | --- | --- |
|  | M (7) |  | F (2) |  | M (12) |  | F (5) |  |
|  | VH. Cells (x10<sup>6</sup>) | Tumor Vol. | VH. Cell (x10<sup>6</sup>) | Tumor Vol. | VH. Cell (x10<sup>6</sup>) | Tumor Vol. | VH. Cell (x10<sup>6</sup>) | Tumor Vol. |
| Before Xenograft | 2.684 | 0 | 1.302 | 0 | 2.507 | 0 | 2.596 | 0 |
| After Xenograft | 2.917 | 10.47 | 1.656 | 1.047 | 2.774 | 7.54 | 2.887 | 25.55 |
| Before 1<sup>st</sup> Treat | 2.621 | 49.35 | 1.587 | 39.173 | x | x | x | x |
| Before 2<sup>nd</sup> Treat | 3.579 | 50.88 | 3.150 | 37.608 | x | x | x | x |
| Before Euthanizing | 2.235 | 45.84 | 2.704 | 85.694 | 2.542 | 94.53 | 2.728 | 149.01 |

\*VH. Cells: Average Cell counts in the vitreous humor, x: data was unavailable.

The average cell count in vitreous humor/density value in healthy mice eye (before xenograft) is not zero for NSGS and Nu/Nu female mice and is found to be almost equal. The number of cells after the xenograft process increased in all cohorts, irrespective of the initial number of cells injected. The cell population decreased between xenografts till the first treatment is applied, and the tumor volume increased. The cell population decreased further between the first and second treatments in the case of the NSGS Female cohort. However, for Nu/Nu, this value increased, showing the only difference. As the multiple treatment sessions are conducted right before euthanizing, the cell population became comparable (never went lower) to values of healthy mice conditions for the female cohort. For the male cohort, the cell population right before euthanizing became lower than it was before xenograft. The tumor volume decreased for the treatment cohort but increased for the cohort that is given no treatment. The next section explores an exhaustive time-independent correlation between gender, genotype, and cell population with tumor volume. Figure 6 shows the mean, max, and minimum values along with overlapped tumor volumes, giving further confidence and the need to look for any possible correlations.

**Figure 6:**
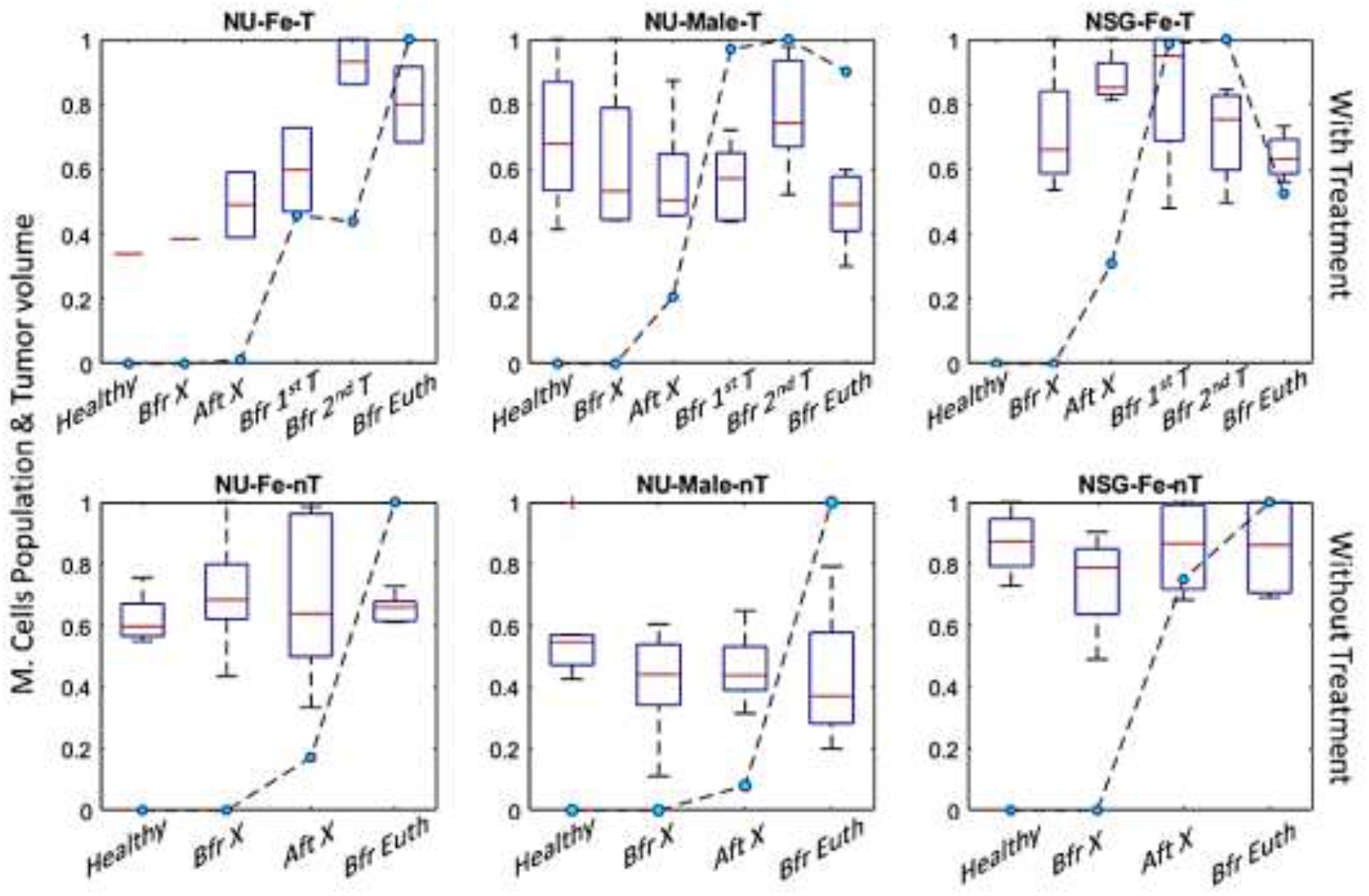
Normalized Average counts Cells in vitreous humor / Population (box plot) and Tumor Volume (plot with blue marker) at different stages (i.e., Healthy, before and after xenograft, before 1^st^ and 2^nd^ treatment, and finally before euthanizing) in Nu / Nu an NSGS female and male mice subjected to treatment and without treatment protocols. The average Cell Population follows the trend in growth in Tumor volume.

### 3.5. Histopathological studies

Histopathology of mice eyes are investigated for confirmatory evidence of the tumor microenvironment *in vivo* and extended biological information. Figure 7 shows the results of the same animal that is used in Figure 5. Selected data where the experiment prolonged for the maximum number of days has been used and compared with the histological data. Figure. 7 represented the qualitative evaluation of invasive and non-invasive imaging, which shows the *in vivo* identification of immune cells due to inflammation with its maximum tumor growth (in the left eye) that grows out of the thin retina.

**Figure 7.**
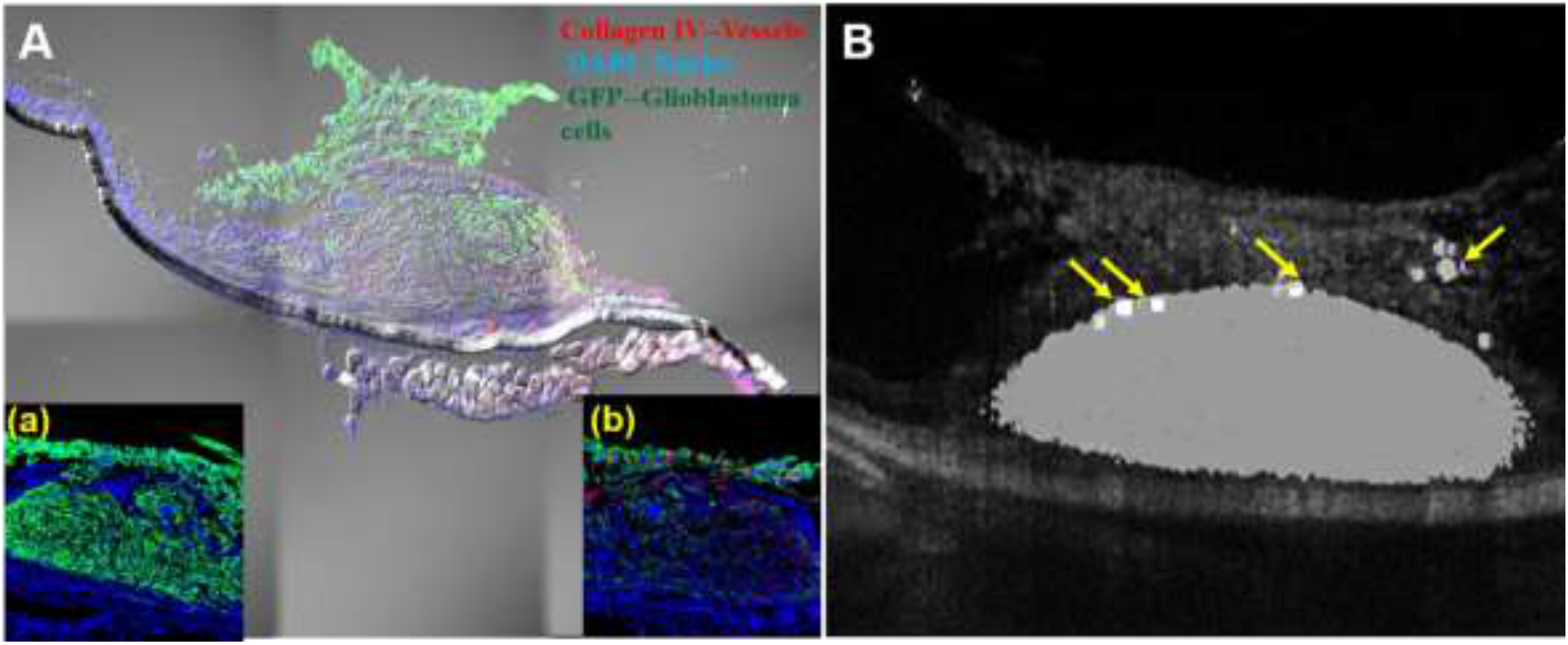
Visualization of immune biomarkers by invasive and non-invasive methods. **A.** Invasive imaging via confocal microscopy: Histopathological analysis of Athymic nude-foxnlnu (Nu/Nu) mice (left eye) showing the infiltration of CD8+ cell surface markers onto tumor and the laser burned retina. The magnified confocal micrographs of dye-stained cellular components are shown in the inset of Figure 6A. Figure 6Aa shows the absence of CD4+ cell surface markers, and Figure 6Ab shows the infiltration of the CD8+ markers confirmed by the flow cytometry analysis. The right eye shows no tumor growth, (Both eyes are sacrificed and performed histology experiments). **B.** Non-invasive imaging of the same left eye using optical coherence tomography (OCT) illustrated the immune cell infiltration indicated by arrow symbols.(*OCT imaging was done before the histological analysis*).

The tumor is immediately identified from the cryosections (Fig. 7(A)) and imaged using confocal microscopy used to assess the changes in the physiological features. Tumor neovascularization indicates the blood vessels intrinsic to the tumor-associated with choroid vasculature. Thus, the neovasculature was corroborated through collagen IV staining (Figure. 7). It also revealed the normal retinal layering and the tumor grown between the retinal tissues tagged with GFP and DAPI fluorescent markers for differentiation. In addition, we also observed the most important features of inflammatory signals of CD8+ cell surface markers (very high immune cells) infiltered onto the tumor cluster of the left eye (Fig. 7Ab). Unfortunately, there are no CD4+ markers found in Figure 7Aa. These are co-receptors for T-cell receptors found in monocytes, macrophages, dendritic cells, and helper T-cells. Besides, the histopathological studies are also correlated with the processed data and support the proposed scientific claim. However, in the case of the right eye, the tumor disappeared due to retinal atrophy, and its histological result is shown in the supplementary section, Figure S8. The invasive imaging result is compared with the non-invasive in vivo image, which confirms the similarity and also observed the immune cells infiltrated onto the bulk tumor cluster illustrated in Fig.7B.

### 3.6. Relation between particle density with xenografted glioblastoma volume

Normalized cell counts in Vitreous humor (VH. Cell) and volume of a tumor with respect to days after xenograft are plotted in Figs.S5 and S6. The inclusive correlation study suggested that the immune cell generation has been influenced by the tumor growth progression, which is associated with and supports the hypothesis. Further, the evaluation was done by separating the data sets into exclusively non-treated ones and the treated data sets. It is to be noted that few xenograft eye models are lately treated using ‘nanodox’ (nanodrug of doxorubicin) during the experiment. The treated data sets are further divided into two regions, before treatment, and after treatment. These two regions explicitly give information on the change in the mathematical expression of the data sets before and after treatment without considering gender and genotype.

#### 3.6.1. Visual Comparison of plots

Alternatively using cross-correlation or simple correlation analysis one can visually observe by plots. Figures S4(c) and S5(c) show vitreous cells and tumor volume w.r.t time.

For Nu/Nu female mice without any treatment protocol, the relationship between cell and tumor volume is proportional for all 7 cases. For 5 out of 7 cases both parameters increased. Nu/Nu female treatment cases are sparse in number and are considered inconclusive here.

For Nu/Nu male mice cohort without any treatment, the relationship is proportional for 8 out of 20 cases, inverse proportional for remaining cases. However, the cases with inverse proportionality could only survive for relatively less duration (less than 25 days). During the treatment this relationship got converted into proportionality as the tumor volume and cell density decreased.

For NSGS female mice without any treatment protocol the relationship between these two quantities is inversely proportional for 5 out of 8 cases, and flat in remaining 3. Where the tumor volume increased around day 20 and suddenly decreased significantly afterwards (even without treatment) in 4 out of those 5 inverse proportional cases, the cells density increased with time in these cases. The observation is reported earlier^64,65^. When treatment is introduced the cell density as well volume, both parameters decreased in 3 out of 4 cases.

#### 3.6.2. Correlation after considering gender and genotype

Collective information is summarized in Figure 8. Boxplot of correlation between cell density values of NSGS, Nu/Nu male and female before and after treatment is shown. The trend shows (Figs. 8A) a weak positive relationship between cell density and tumor growth for Female Nu/Nu with a median correlation value of 0.3. A weak correlation with a value of −0.29 is estimated for NSGS female mice. For male Nu/Nu mice, this relationship is even weaker, having a median correlation value of −0.18 (Fig. 8C). The treatment, however, has inverted the depicted relation for all three categories towards mild-strong relations. The Female NSGS and Male Nu/Nu mice cohort show a positive correlation with 0.63 and 0.5 median values, respectively. The data, although, shows significant variation. It may own to the fact that all these mice are injected with different amounts of tumor cells. Weirdly, but for Female Nu/Nu mice (only 2 cases combined), the median correlation value effectively becomes −0.53.

**Figure 8:**
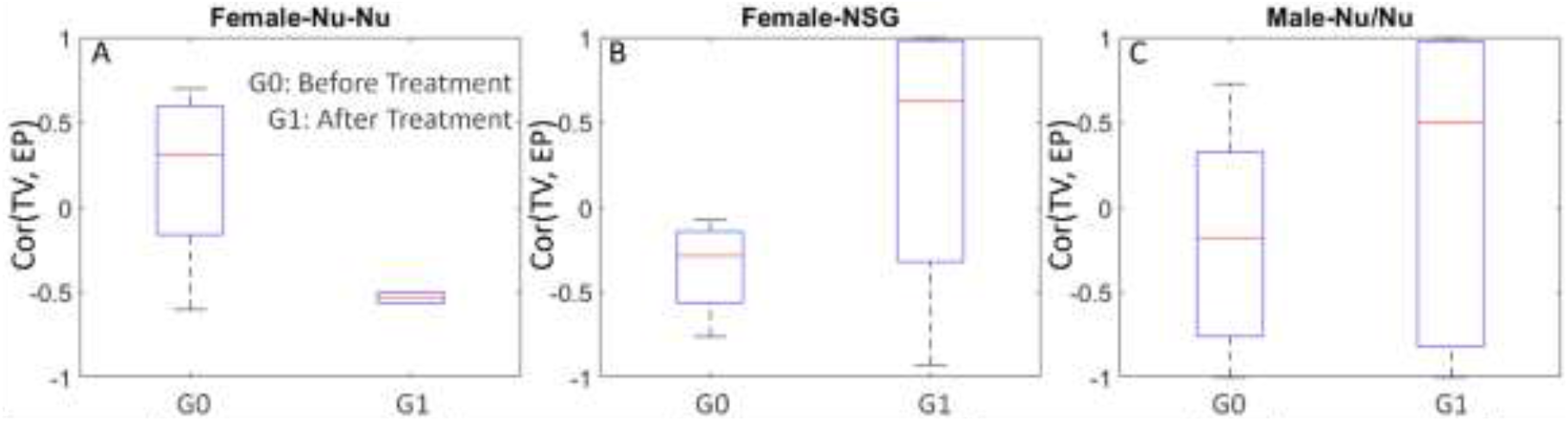
Correlation (overall average including all mice of same genotype) between cell density value and tumor volume; **A**. shows Female Nu/Nu cases before, and after treatment, **B.** shows female NSGS mouse data, and **C.** shows male mouse data.

#### 3.6.3. Cross-Correlation

A numerical investigation provides a cross-correlation between immunosurveillance factors and with tumor growth model^66^. Cross-correlation values are estimated to measure the similarity and coherence along with time-dependency between normalized tumor volume growth and normalized vitreous cell density.

Figure 9 depicts these cross-correlation values (x-cor values) plotted with respect to the number of days from beginning to end. The first, second, and third row show the data of Nu/Nu Female, Nu/Nu Male, and NSGS Female, respectively. First two columns in each row (Figs. 9(a), (b), (d), (e), (g), and (h)) belongs to cases without treatment. The last column (Figs. 9(c), (f), and (i)) is of treated cases. The fitted curve to the x-cor values is plotted in red dot marker along with original values in blue dot marker. The shaded region depicts an error in the fitting. The figures are categorized according to the optimal degrees/order (linear/1^st^, quadratic/2^nd,^ and cubic/3^rd^) of the fitted polynomial. The curve fitting using a higher degree often leads to the approximation or generalized results lacking the true characteristics of the functions and hence avoided. The first column contains data that has the least root mean square error (as a goodness of fit) for 2^nd^ order polynomial termed genotype-gender-quadratic-nT. Similarly, the second column contains the third order. Only Nu/Nu males have few data fitted best in first-order linear polynomials (shown in the supplementary file as Figure S7). None of the data fitted better for a higher degree than three. The last column, data (treated cases), has non-continued graphs. Although these are from the same animal, but break signifies the start of the treatment protocol. The graph in green contains data till the tumor was allowed to grow without any treatment. Each graph is indexed using two indices. The first index refers to the serial/tag or identification number of the respective genotype. The second index refers to the degree of the polynomial. The indexed scheme is used to illustrate that after treatment order of fit of cross-correlation between vitreous humor cell density and tumor volume may change. For example, Fig 9(f) plot tagged as the 2^nd^ order (6,2) transformed into a linear relationship (6,1). Male Nu/Nu untreated, including right before treatment cross-correlation, have eight mice data, all second-order polynomial fits, and nine third-order polynomial fits. It indicates that as the mouse aged and the days progressed after xenograft, for 8 (second-order polynomial cases) mice, the tumor volume and with the number of vitreous cells followed the same non-linear pattern (inverted parabola). This point can be verified by Fig. S5 (b8). Observing the inverted parabolic graphs in the figure below, it appears that the rate of cross-correlation peaks around 75 days, irrespective of the genotype and gender. Similarly, in the rest of the parabolic cases, this rate shoots around 50 days. The treatment effect shows transforming cases having second-order cross-correlation relation into linear relation for Nu/Nu females if introduced fairly late (~100 days); otherwise, only the pattern changed from upward parabolic into an inverted downwards parabolic pattern keeping the degree same. Close observation of Figs 9, S5, and S6 reveals that cross-correlation value fit of 1^st^ order has inverse proportionality, and for higher order, this relation is proportional in nature. The model transformation is found 100% in the case of Nu/Nu Female (however, data is relatively sparse as only two cases are presented) than in the case of Nu/Nu Male (2 out of 5). In the case of NSGS, this transformation is not seen in any of the 4 cases.

**Figure 9.**
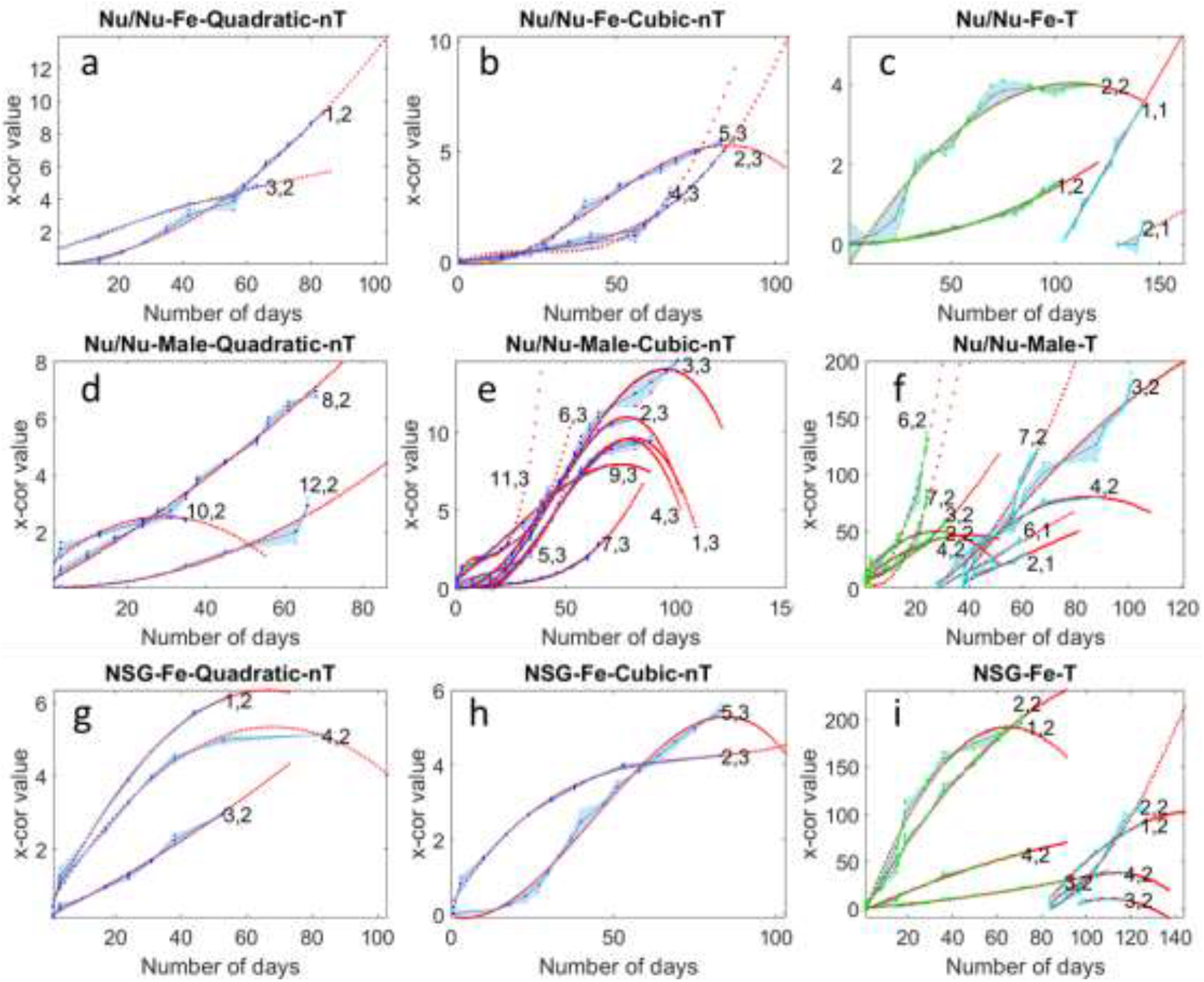
Cross-correlation values between normalized tumor volume and normalized vitreous cell density with respect to the number of days after xenograft. The blue dots represent cross-correlation values, and the red dots represent the fitted curve. Figs. 9(a)-9(c) is Nu/Nu Female; Figs. 9(d)-9(f) Nu/Nu Male; Figs. 9(g)-9(i) is NSGS Female data is used. The third column depicts data before treatment and after treatment. Each graph is indexed with a cage tag of the mouse and the order/degree of the polynomial used to fit the data. After treatment degree of polynomial decreases if treatment starts with a certain delay. The shaded region represents the error in the fitting.

## 4. Discussion

The immunoglobulins test revealed that IgG and IgM scores of female Nu/Nu nude mice remain slightly higher than male Nu/Nu nude mice ^67^. Haematology results show that female Nu/Nu mouse has more Lymphocytes (αβTCR and other) than male Nu/Nu mouse as age progresses^68^. As compared to the Nu/Nu nude mouse, the NSGS mouse does not have B Cells. While innate immunity remains impaired in both genotypes, *NK activity in NSGS remains impaired*, whereas, in Nu/Nu, NK cell density increases with age^69,70^. We note that these findings are not specifically estimated for vitreous humour. Immunophenotypic analysis showed that 10 micron hyalocytes in rat vitreous has tissue macrophages function^71^. Post-chemotherapy test such as Granulocyte-colony stimulating factor (GCSF) treatment increased the number of nonclassical and blood monocytes, and neutrophils in NSGS mice, however does not provide the best environment for the generation of human myeloid cells ^48^. It is also shown that doxorubicin eliminates the myeloid-derived suppressor cells and enhances the capacity of B Cells to activate T cells by enhancing the CD8+ T-cell proliferation and CD4+ Cells responses, which were found in histopathology analysis of the our data set^72^. Tumor growth depends upon effector (for example NK cells) to suppressor cell ratios (including myeloid cells)^73^. At this point, we would like to recall the observation of significant natural presence of the cells, perhaps hyalocytes, in healthy retinal conditions. The already reported evidence of their function include regulation of the vitreous cavity immunology, and modulation of inflammation (74). These last two observations inspire to investigate hyalocytes function as immunosuppressor cells. Due to dynamic nature of this study, it could be mix of cell types. Overall, the presented analysis indicates that, most likely, the observed vitreous body cells are effectively, immunosuppressor cells (Treg or myeloid-derived suppressor cells), as reported earlier^7,41^.

In brief, it is safe to say that an explicit shift in the order/polarity of the model after treatment is corroborated by the affected immune response. In addition, the nanodrug treatment further provides a synergistic immune response along with the innate response. In a nutshell, the mathematical correlation and cross-correlation between the cell growth pattern and the tumor volume provide quantitative and significant biological information on xenograft glioblastoma treatment. This critical evaluation may assist biomedical scientists/physicians in the early diagnosis of any carcinoma and devise therapeutic strategies for treatment by the visualization of these cells in the vitreous humor.

## 5. Conclusions

The present work utilizes automated AI-based segmentation methods to estimate vitreous body cells above the nerve fiber layer (NFL), which facilitated their 3D co-localization with tumor, along with mathematical cross-correlation and correlation. Further, the cell dynamics along the tumor are also depicted, and its infiltration into the tumor cluster is also co-localized using 2D and 3D ocular constructs. In addition, the processed data has been compared with the histopathology analysis for confirmatory evidence. As the tumor volume and vitreous body cells density is shown to be mutually cross correlated, it is safe to say that this relation would represent effector to suppressor cell ratio as well. It thus can be considered as a biomarker for tracking cancer growth in early stages especially owning to the fact that these cells are present in vitreous humor before xenograft.

This work is a combination of cancer biology and the application of programming for the visualization and quantification of immune cells. In a nutshell, this work developed a generalized rapid method that could be an appropriate vitreous biomarker for early diagnosis of inflammation-related disease and ocular cancer.

## 6. Declarations

### Ethics approval and consent to participate

Animals were cared for and handled in accordance with National Institutes of Health Guidelines for the care and use of experimental animals and protocols approved by the Institutional Animal Care and Use Committee of the University of California, Davis between Feb. 2015 till August 2016.

### Consent for publication

Every person has given permission to use their images in this article.

### Availability of data and materials

The data will be made available on request.

## Acknowledgments

MG is thankful to the Department of Science and Technology-Science & Engineering Research Board (DST-SERB) under IMPRINT 2 scheme, Project number: IMP/2018/001045, Government of India. MG is thankful to Dr. Rajkumar Sadasivam, and Dr. Khushboo Gulati for having discussions. MG is also thankful to EyePod, UC Davis for reanalyse the data set.

## CRediT authorship contribution statement

MG: Methodology, Data Acquisition, Investigation, Software, Writing, Visualization, funding; SS: Investigation, Software, HW: Histopathology, PZ: Data Acquisition, RZ: Investigation, funding.

## 8. Supplementary

### 8.1 Vitreous Cells before xenograft in healthy mouse eye

**Figure S1:**
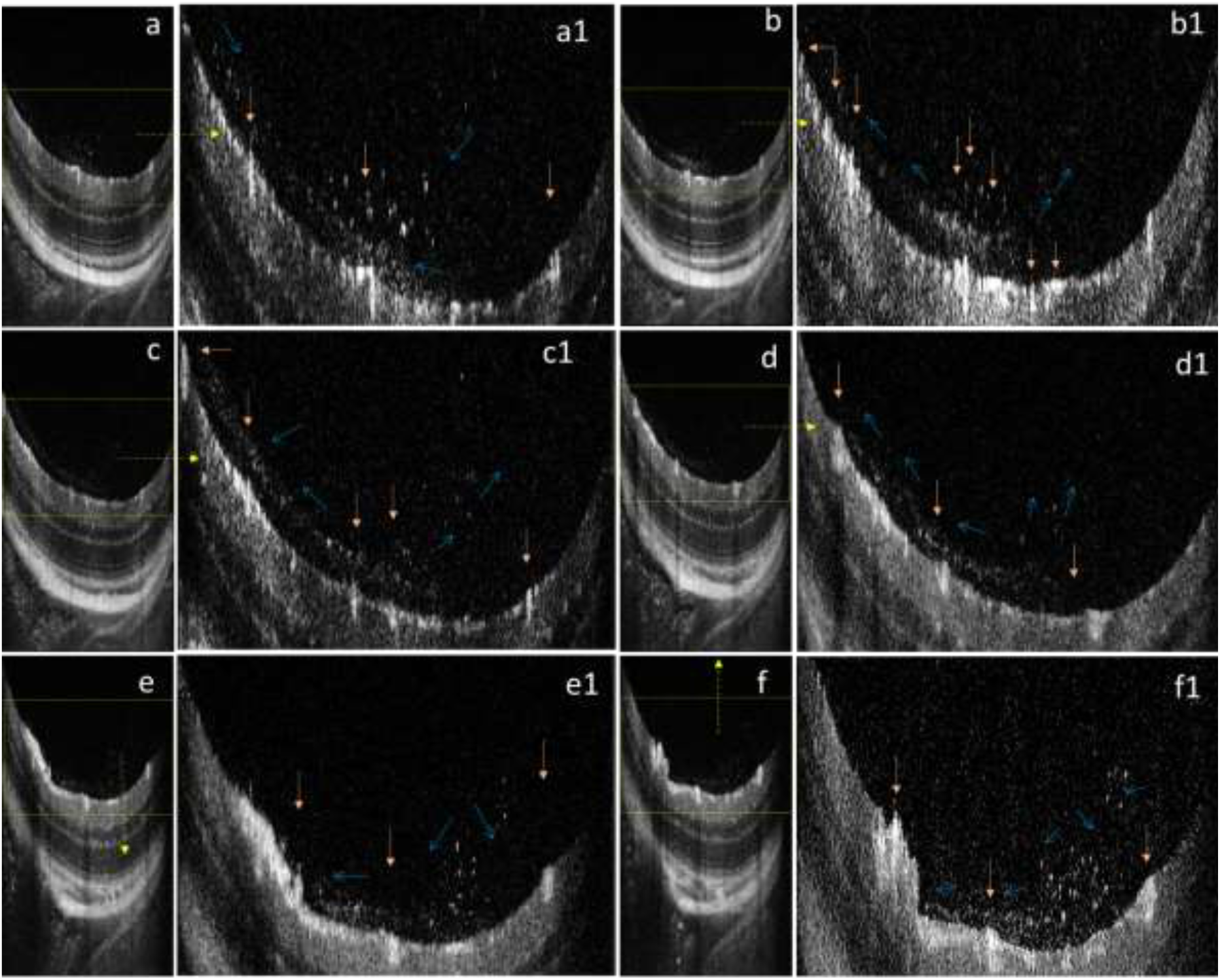
Presence of Vitreous humor cells in healthy Nu/Nu Female Mouse *<u>before Xenograft</u>* in different positions. Figs. (a) – (f) are Bscans and (a1) – (f1) are respective portion containing ILM till Vitreous humor body (with necessary digital zoom). Figs (a1), (b1) shows relatively higher cell density to the proximity of blood vessels (marked with orange color vertical and horizontal arrows), may be acting as a source of these cells. These cells in form of stream (marked by blue thin arrows), are migratory in nature, stay near to the ILM. The migration w.r.t blood vessels indicates communication between several blood vessels via vitreous humor. Please refer to <u>Movie M0</u> for full data.

### 8.2 Stream of cells between the optical cord and other blood vessels

**Figure S2.**
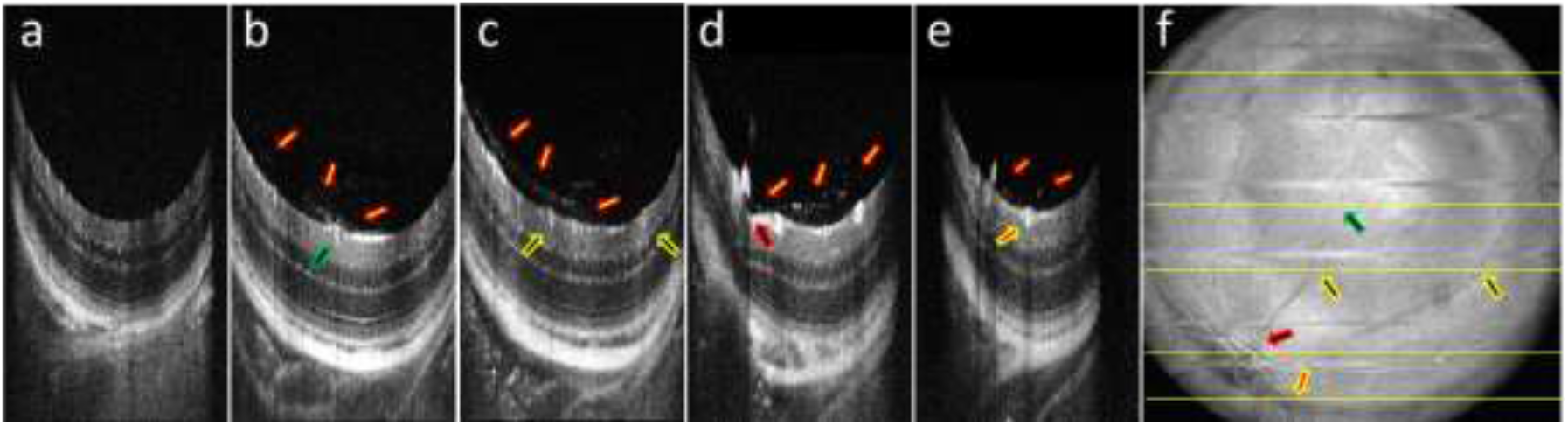
Presence of cells in Vitreous Humor of Nu/Nu Mice Female *before* Xenograft. Figs. S2 (a-e) shows B-Scan marked from top to bottom in En-Face shown in Fig. S2 (f). Yellow Arrows with red boundaries (in vitreous humor) shows the stream of cells in each image. Rest of the arrows (inside retina) shows origin of stream near to the blood vessels. These second set of arrows are marked in Enface with corresponding locations. <u>Movie M2</u> shows full healthy retinal OCT data. This composite image shows that stream of cells are connected between optical cord and other blood vessels near to ILM inside vitreous cortex only.

### 8.3 Capability of imaging technique and cells in 3D

**Figure S3.**
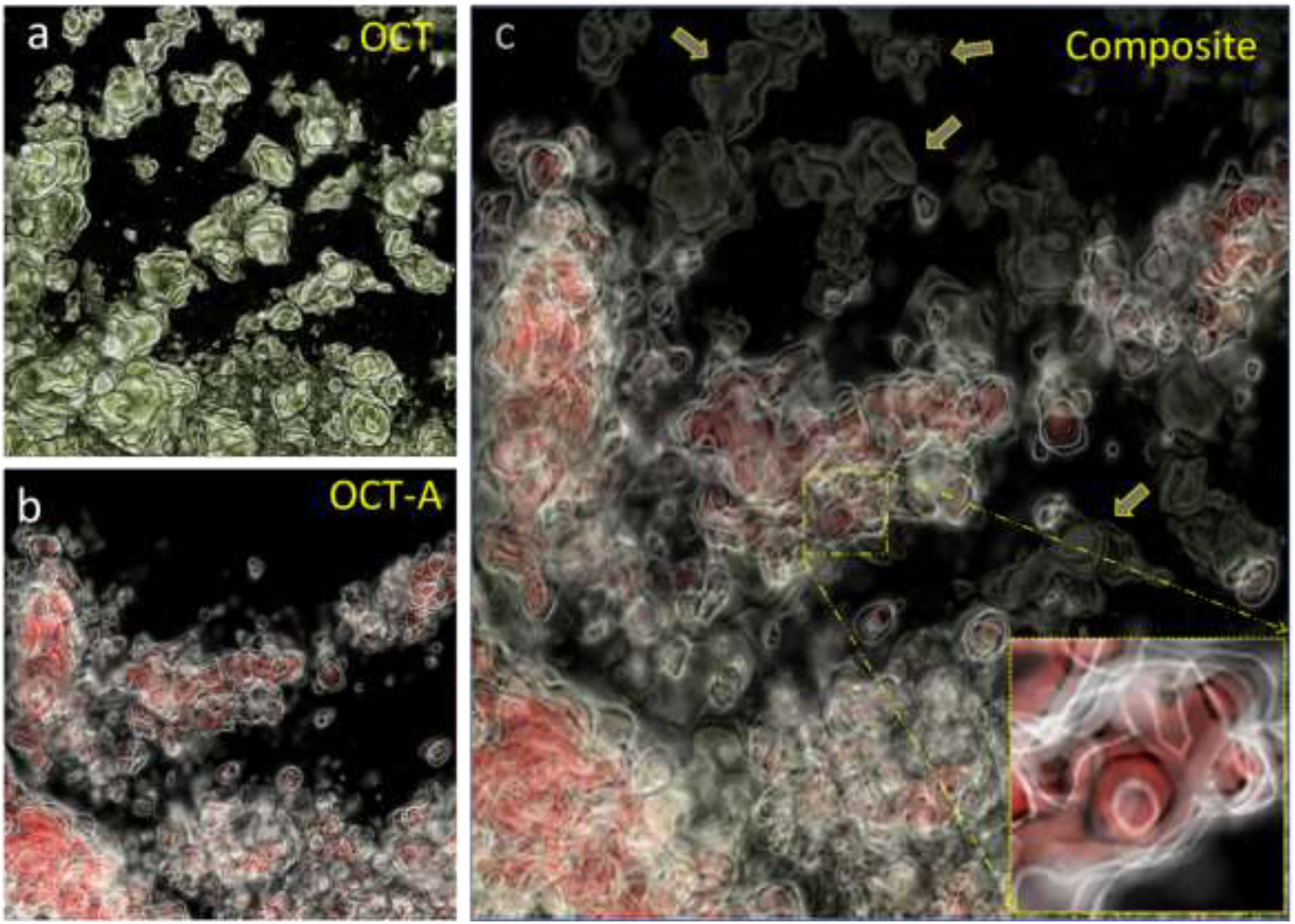
Composite images highlight the presence and absence of structures relative to OCT (artificial green shade in this image) and OCT-A images, taken simultaneously. Figs. S3(a) and S3(b) slightly differ in projection angle. It indicates that specific structures missing in OCT-A were not experiencing a phase change, possibly due to negligible movement. The red part refers to the portion inside the blood vessel with motion. These structures (visible in OCT but not in OCT-A and marked using yellow arrows) Fig. S3(a) have discrete structures indicating they are not part of the retina but clumped cells. The donut shaped blood cell (~7 micron) shown in inset image in Fig. S3(c) shows capability of imaging technique providing a scale of immunoglobulin cells in vitreous humor.

### 8.4 Dynamics and infiltration of immune cells

Three dimensional (3D) images of the retina were constructed to evaluate immune cell migration, its infiltration, and tumor growth pattern (Figs. S4. (B) and S4. (c)). The immune cells could be explicitly seen in the timeline 3D structures represented in column 2 of Figure S4. (B). Subsequently, the dynamics of the immune cells can be identified and visualized in 3^rd^ column of Figure S4. (C). In brief, the timeline 3D images of Figure S4. (C) (column 3) clearly indicated that on 1^st^ cycle of imaging, the image was sliced at a certain distance from optic nerve, whereas during the 9^th^ cycle, the image segmentation was done at a different slice. Similarly, the slicing varies because the location of the distribution of cells varies with time due to kinetic motion and the pursuit of obtaining the exact morphology of cells.

This attributed that the location of the particles varies with the timeline images, which further explained that immune cells are dynamic in nature. The above findings confirmed the migration of immune cells along with the tumor growth. Figure. S4. (c) represented the 3D sliced images of the immune cells (as elusive particles) showing the migration and infiltration into the tumor over the timeline of random events that occurred in between 1 to 150 days. The 3D sliced images has been compared with OCT B-scan 2D images of Figure S4(D) indicated the cross-section of the tumor with precise localization of infiltrating immune cells through the XY plane. Also, the mapping analysis was done to visualize the immune cell generation and its respective tumor behavioral pattern in the microenvironment. The mapping was shown in *en face* images with different color schemes, i.e., dark red (immune cells), yellow (tumor growth), and the rest of the layers (grey). The 3D constructed scheme for visualization exclusively illustrated the immune cell generation, migration, and its corresponding tumor growth. The constructs are represented in uniform angles for a clear vision of both immune cells and tumors. This result suggested that the immune cells can be identified as prognostic markers that initiate tumor growth, which could be used as a diagnostic biomarker for cancer prediction.

#### 8.4.1. 3D Visualization and Co-localization of immune cells

Multimodal imaging system OCT and SLO have been used for the *in vivo* investigation. The acquired data are processed to evaluate the relationship between tumor growth pattern and the presence of myeloid cells in the vitreous humor. Figure. S4. illustrated the timeline series of the selective processed data set showing the ocular structure in different perspectives. Figure. S4. (A) represented the *en face* real-time SLO images from 1^st^ cycle to 28^th^ cycle that are color differentiated to identify the immune cells through the longitudinal section (aerial view). It consists of tumor vasculature, choroid structure, and retinal vasculature. The tumor cluster was co-localized using SLO GFP fluorescence superimposed into the choroid. The *en face* images (First column) are overlaid with ocular components such as elusive particles (yellow), choroid, tumor blood vessels, retinal vasculature, and the nerve fiber layer (NFL). Subsequently, the processed OCT data are used to obtain 3D reconstructed greyscale images for visualization of the immune cells and its infiltration into the tumor cluster (shown in 2^nd^ column of Figure. S4. (B).

**Figure S4.**
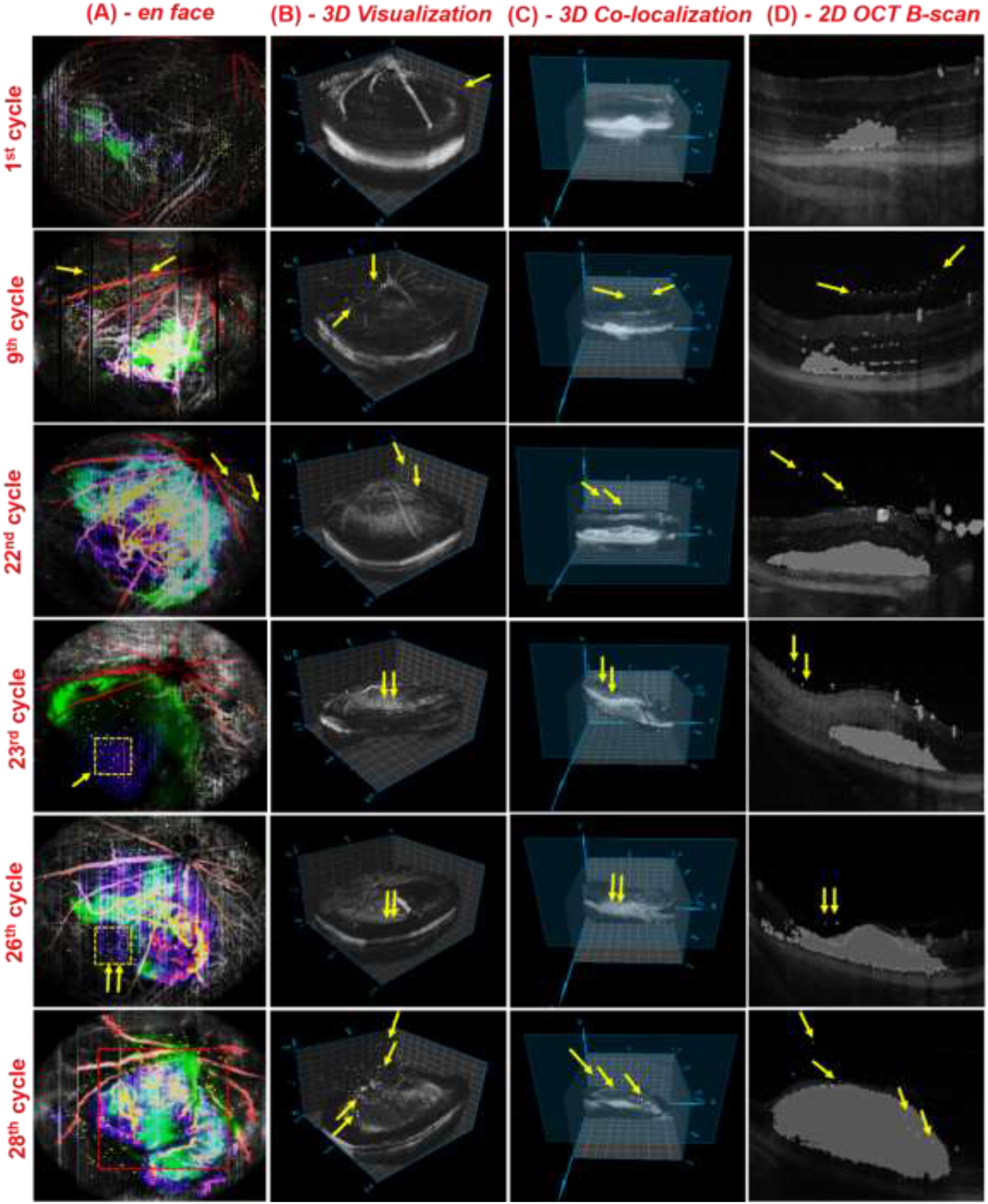
Representation of processed data set showing 3D Visualization and Co-localization of cells. A. Overlay SLO *en face* images illustrating the color differentiated choroid, tumor vasculature, elusive particles and nerve fiber layer (NFL), **B.** Timeline visualization of 3D constructed retinal structure with tumor growth progression along with cell generation (indicated in white coloured particles in column B), **C.** Co-localization of immune cell dynamics in 3-D retinal structure segmented at different Z-slices shown in Figure S4. C. (Each cycle represents the particular day in which the OCT imaging was carried out during the study) **D.** XY 2D B-scan OCT images showing the cell generation and dynamics in tumor microenvironment. (The cell dynamics is indicated through yellow arrow marks in Figure S4. (D)).

### 8.5 EP or VH. Cells vs. Days without treatment

**Figure S5.**
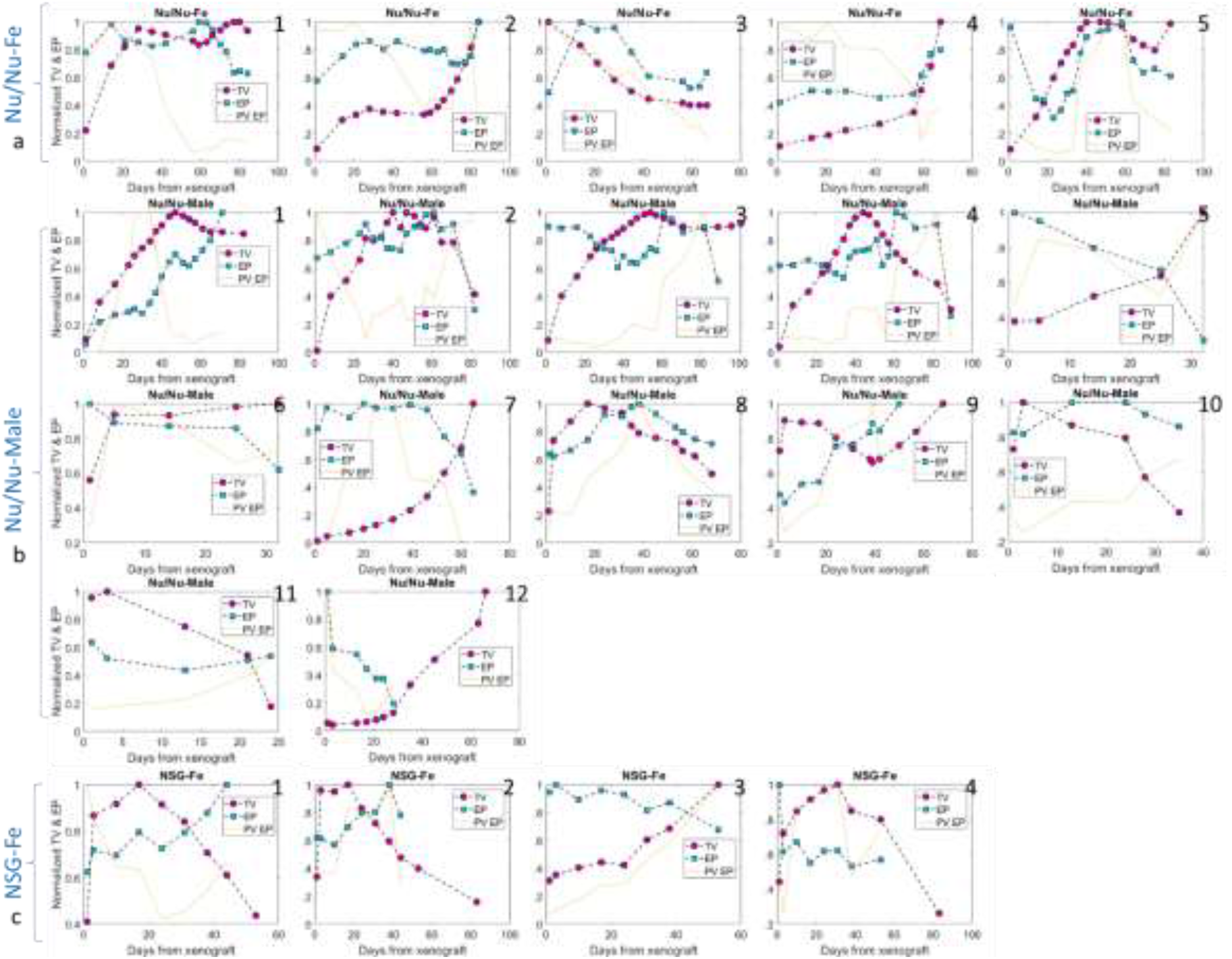
Total Volume (TV) and Average Particle Density (EP or VH. Cells) vs. Days from xenograft plot. Fig. S5(a) in first row, Fig. S4(b) in center rows and Fig. S5(c) in last rows contains data plots of Nu/Nu Female, Nu/Nu Male and NSG Female mice cohort, respectively. None of the mouse in this figure is treated. Most of the graphs show almost a linear relation being followed between both of the parameters with few exceptions(2,10,12, 15, 18). These exceptions follow inverse linear relationship.

### 8.6 EP or VH. Cells vs. Days with treatment

**Figure S6.**
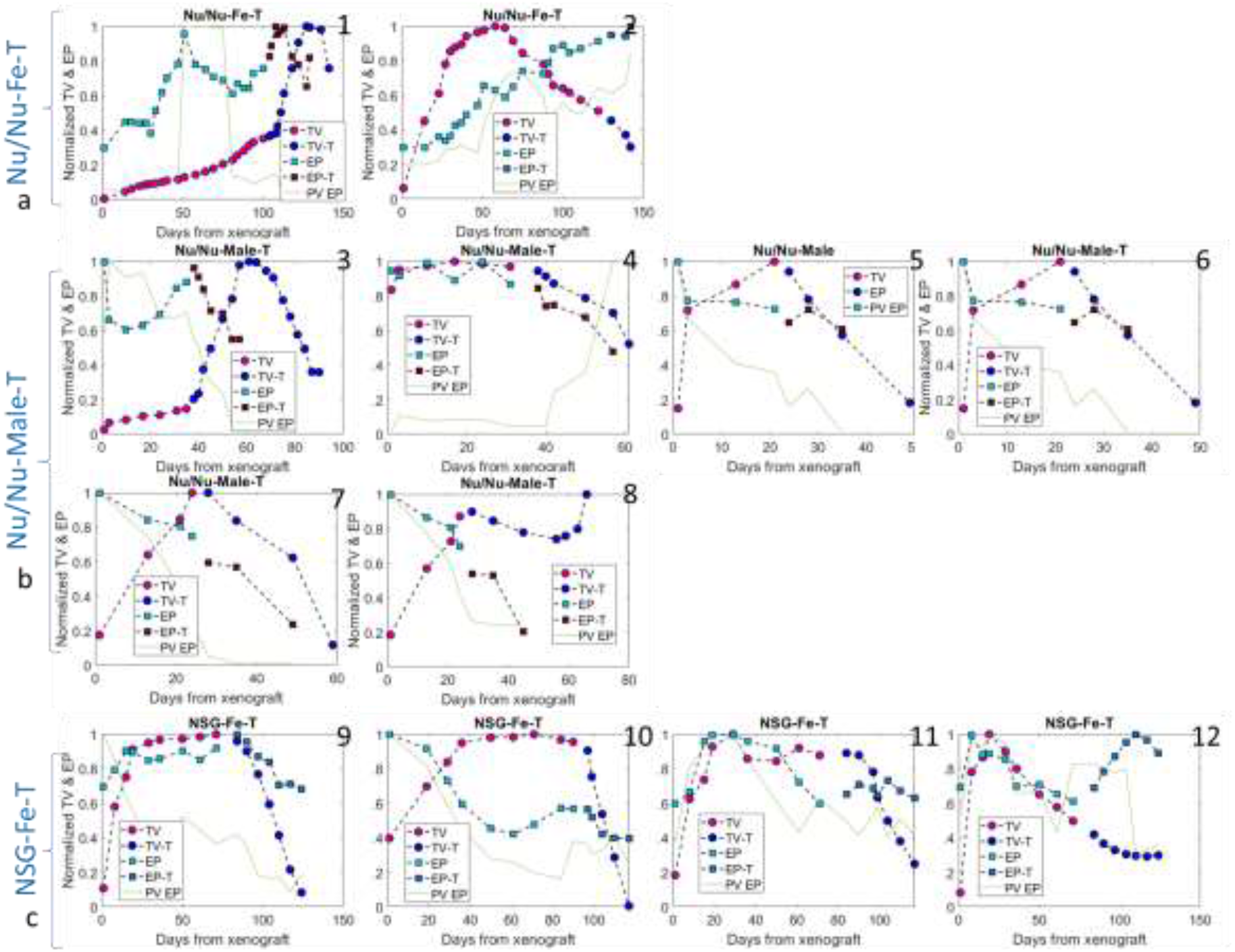
Total Volume (TV) and Average Particle Density (EP or VH. Cells) vs. Days from xenograft plot. Fig. S6(a) in first row, Fig. S6(b) in center rows and Fig. S6(c) in last rows contains data plots of Nu/Nu Female, Nu/Nu Male and NSG Female mice cohort, respectively. All of the mice in this figure have gone under nanodox dynamic photodynamic imaging assisted treatment protocol. The color scheme for marker changes once the treatment data is plotted. Before treatment again almost a linear relation seems to exists between both of the parameters with few exceptions(8,10). After treatment inverse linear relatin is shown in few cases (1, and 12).

### 8.7 Cases with single degree Cross correlation fit

**Figure S7.**
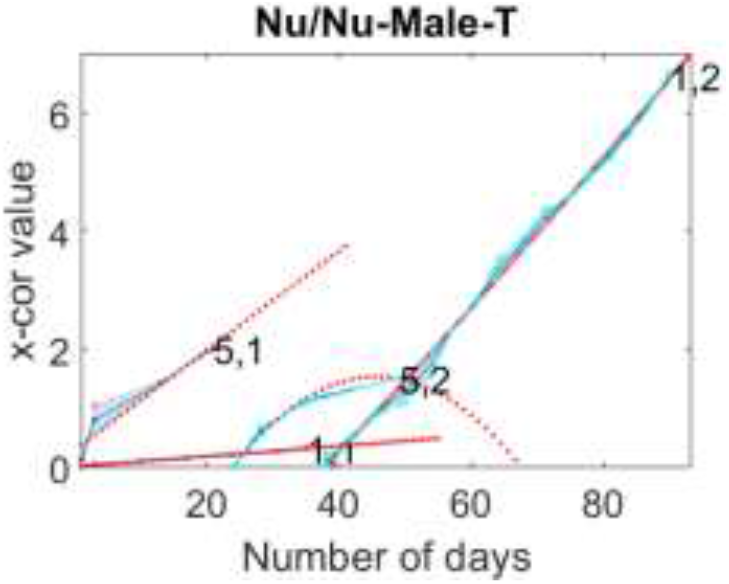
Cross correlation values between number of vitreous humor cells and tumor volume growth fit to polynomial of single degree (linear fit). The red markers are fitted values, dots in cyne color are original values and blue shade represent the error in fitting. The data belongs to Nu/Nu male mouse those have undergone the treatment protocol.

### 8.8 Histopathology showing cells

**Figure S8.**
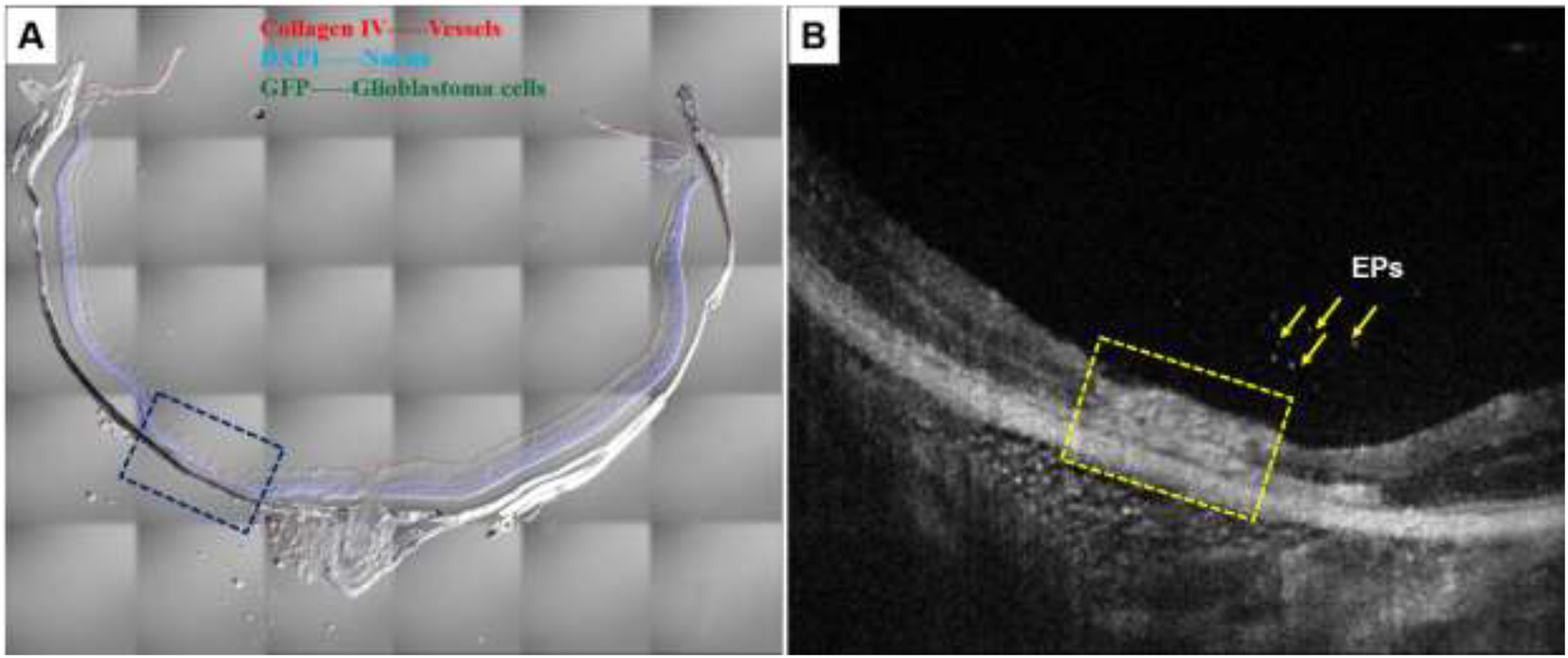
Qualitative analysis of invasive and non-invasive imaging. Invasive confocal microscopic analysis for studying immunohistochemistry. **A.** Athymic nude-foxnlnu mice (right eye), concentration - 250 cells/0.25 μl, with Nanoparticles Treatment (Nanodoxorubicin) and sacrificed on 05/11/2016. After treatment the tumor disappeared with retina atropy (region marked in dark blue). and Non-invasive OCT imaging of **B**. 2D B-scan image of Athymic nude –foxnlnu mice showing the visualization of cells in arrow and the condition of retinal atrophy in XY plane.

**Table ST1:**
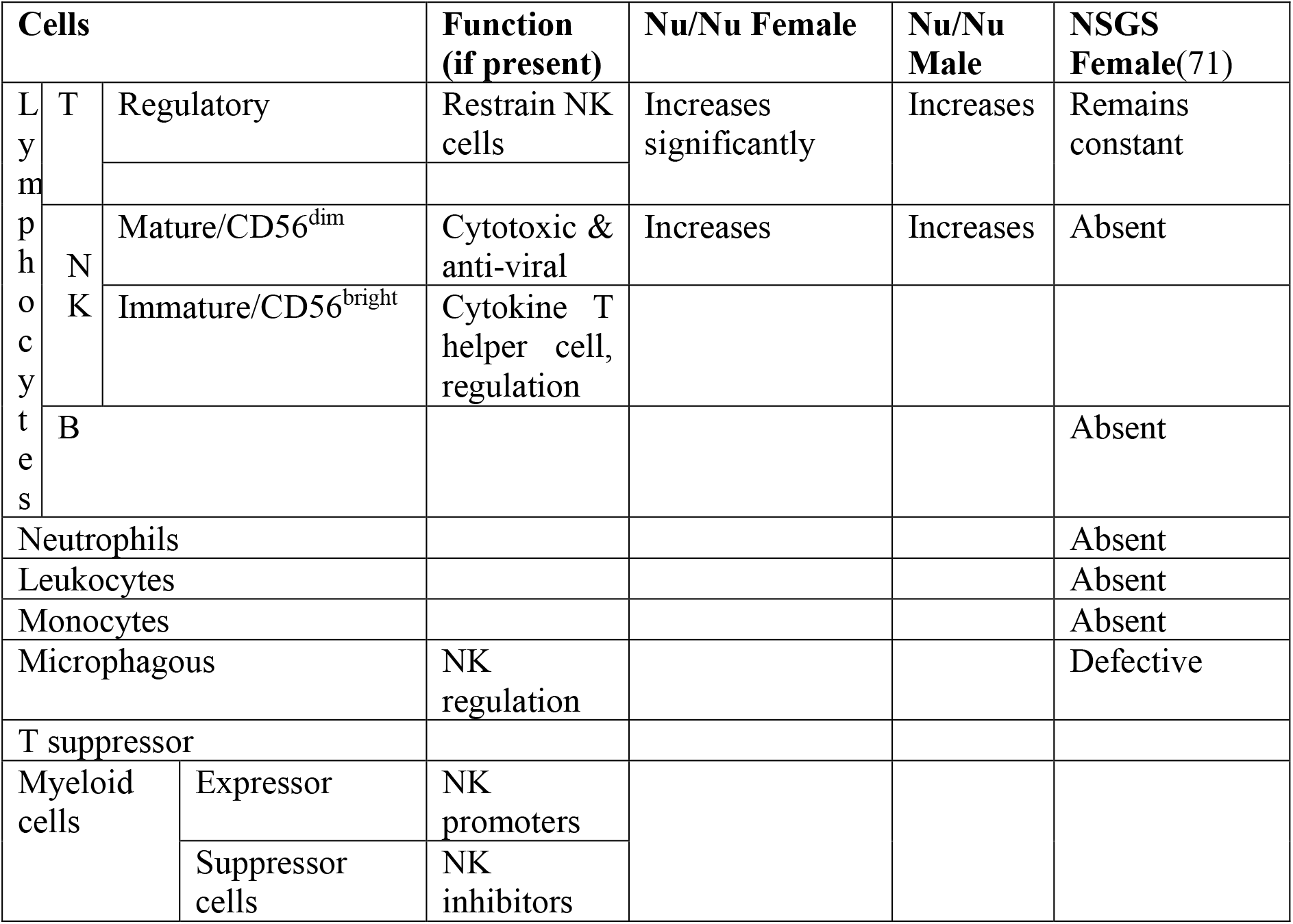
Cells growth w.r.t. age/time.

