## Supplementary material for "Prolonged *in-vivo* tracking of vitreous fluid and its early diagnostic imaging biomarkers for cancer growth": Figure S

### 1. Supplementary

#### 8.1 Vitreous Cells before xenograft in healthy mouse eye

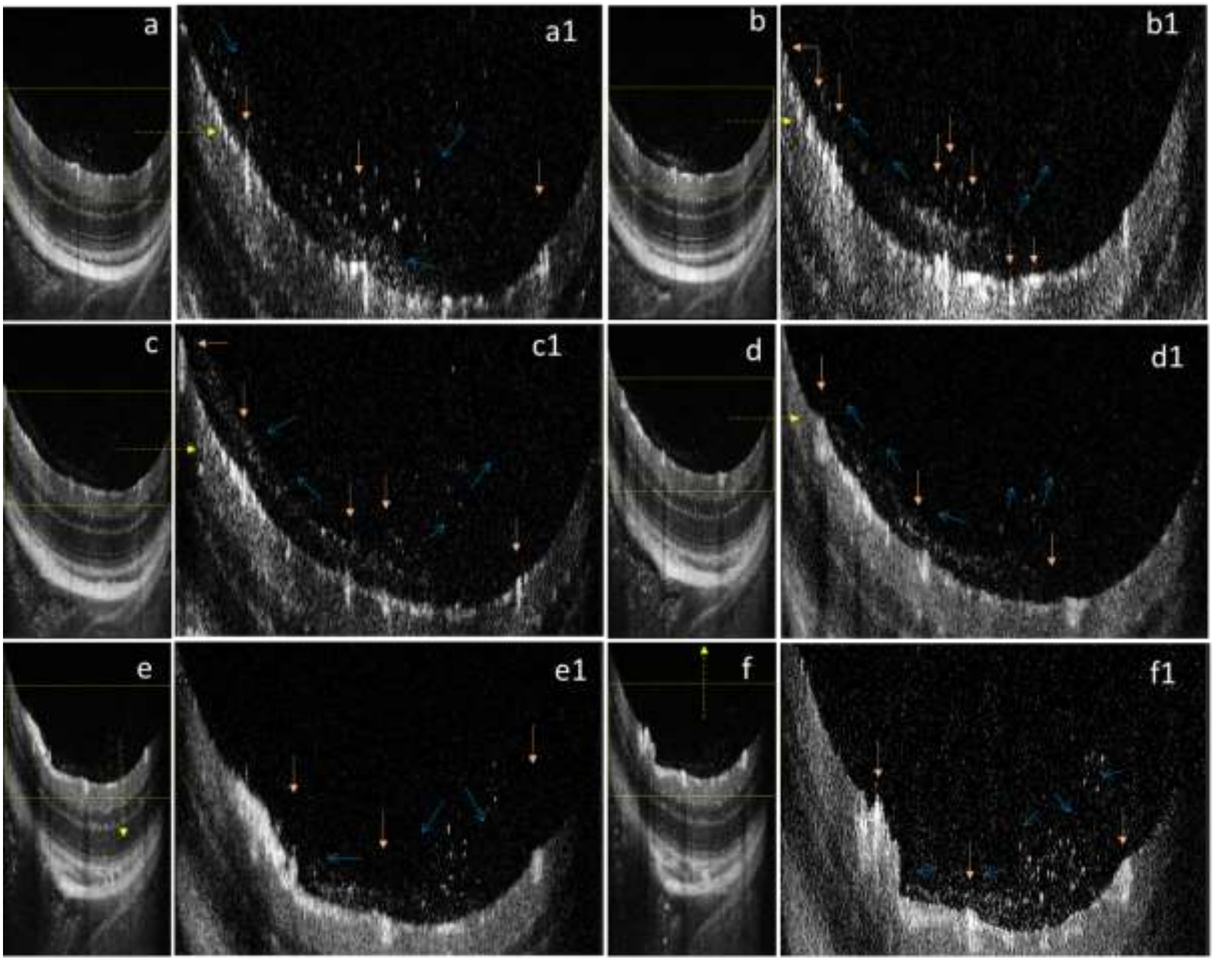


**Figure S1**: Presence of Vitreous humor cells in healthy Nu/Nu Female Mouse *before Xenograft* in different positions. Figs. (a) – (f) are Bscans and (a1) – (f1) are respective portion containing ILM till Vitreous humor body (with necessary digital zoom). Figs (a1), (b1) shows relatively higher cell density to the proximity of blood vessels (marked with orange color vertical and horizontal arrows), may be acting as a source of these cells. These cells in form of stream (marked by blue thin arrows), are migratory in nature, stay near to the ILM. The migration w.r.t blood vessels indicates communication between several blood vessels via vitreous humor. Please refer to Movie M0 for full data.

#### 8.2 Stream of cells between the optical cord and other blood vessels

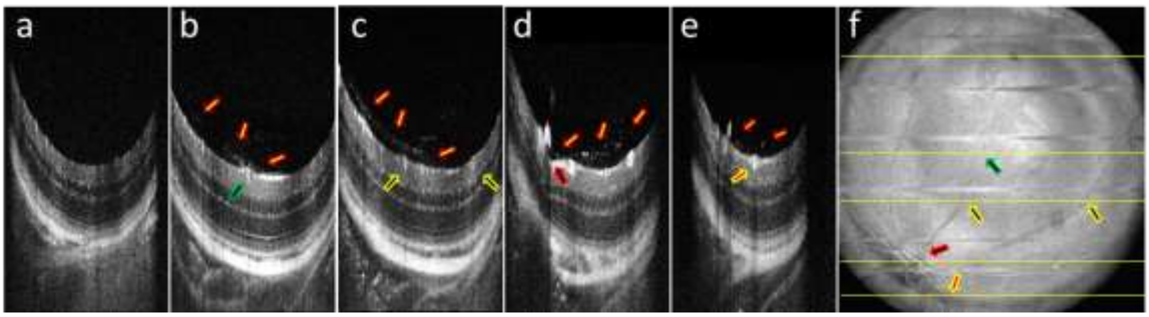


**Figure S2.** Presence of cells in Vitreous Humor of Nu/Nu Mice Female *before* Xenograft. Figs. S2 (a-e) shows B-Scan marked from top to bottom in En-Face shown in Fig. S2 (f). Yellow Arrows with red boundaries (in vitreous humor) shows the stream of cells in each image. Rest of the arrows (inside retina) shows origin of stream near to the blood vessels. These second set of arrows are marked in Enface with corresponding locations. Movie M2 shows full healthy retinal OCT data. This composite image shows that stream of cells are connected between optical cord and other blood vessels near to ILM inside vitreous cortex only.

#### 8.3 Capability of imaging technique and cells in 3D

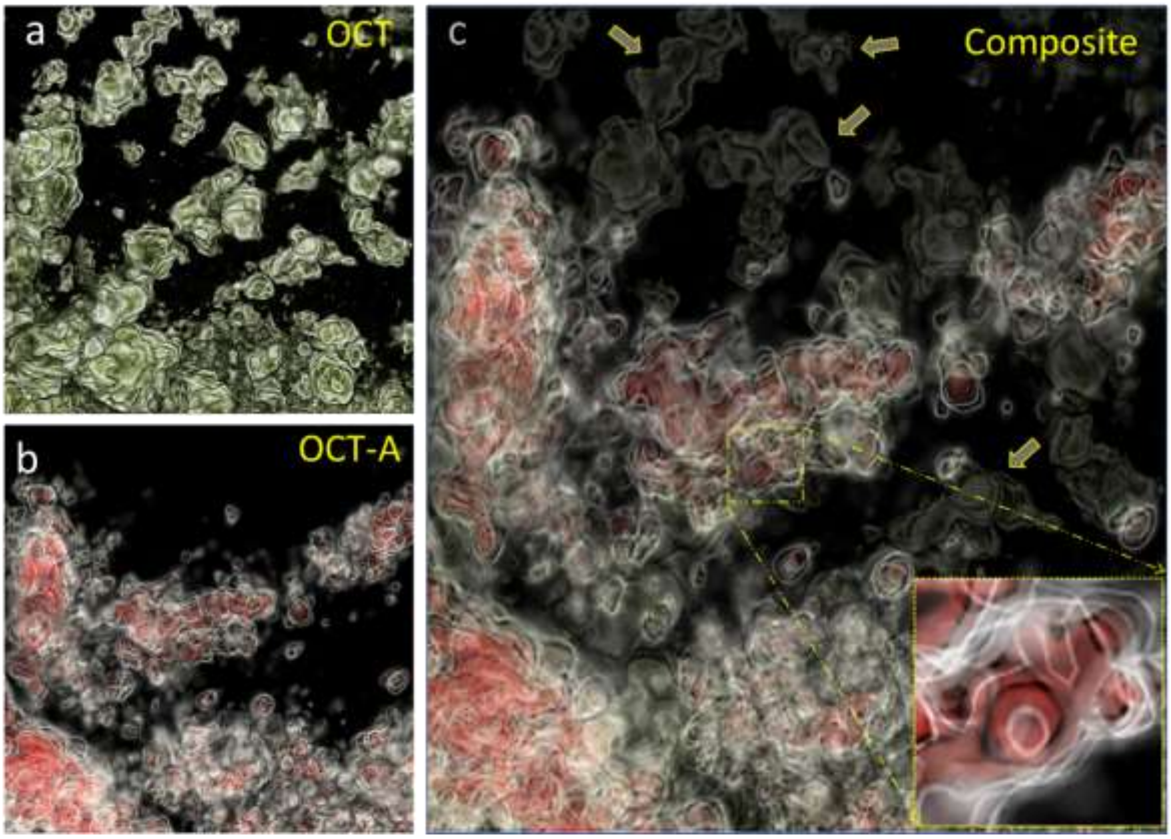


**Figure S3.** Composite images highlight the presence and absence of structures relative to OCT (artificial green shade in this image) and OCT-A images, taken simultaneously. Figs. S3(a) and S3(b) slightly differ in projection angle. It indicates that specific structures missing in OCT-A were not experiencing a phase change, possibly due to negligible movement. The red part refers to the portion inside the blood vessel with motion. These structures (visible in OCT but not in OCT-A and marked using yellow arrows) Fig. S3(a) have discrete structures indicating they are not part of the retina but clumped cells. The donut shaped blood cell (~7 micron) shown in inset image in Fig. S3(c) shows capability of imaging technique providing a scale of immunoglobulin cells in vitreous humor.

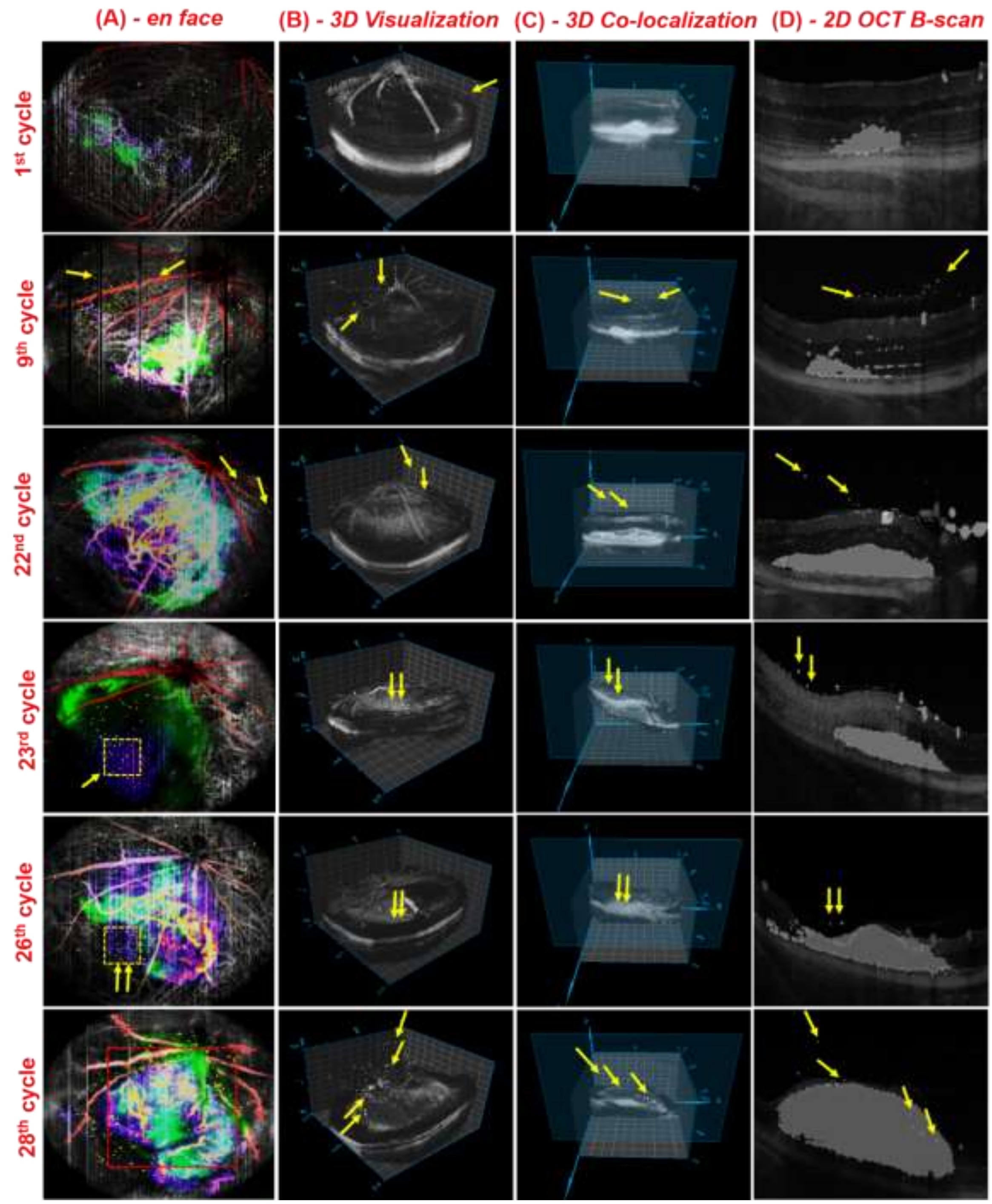


**Figure S4**. Representation of processed data set showing 3D Visualization and Co-localization of cells. A. Overlay SLO *en face* images illustrating the color differentiated choroid, tumor vasculature, elusive particles and nerve fiber layer (NFL), **B.** Timeline visualization of 3D constructed retinal structure with tumor growth progression along with cell generation (indicated in white coloured particles in column B), **C.** Co-localization of immune cell dynamics in 3-D retinal structure segmented at different Z-slices shown in Figure S4. C. (Each cycle represents the particular day in which the OCT imaging was carried out during the study) **D.** XY 2D B-scan OCT images showing the cell generation and dynamics in tumor microenvironment. (The cell dynamics is indicated through yellow arrow marks in Figure S4. (D)).

#### 8.5 EP or VH. Cells vs. Days without treatment

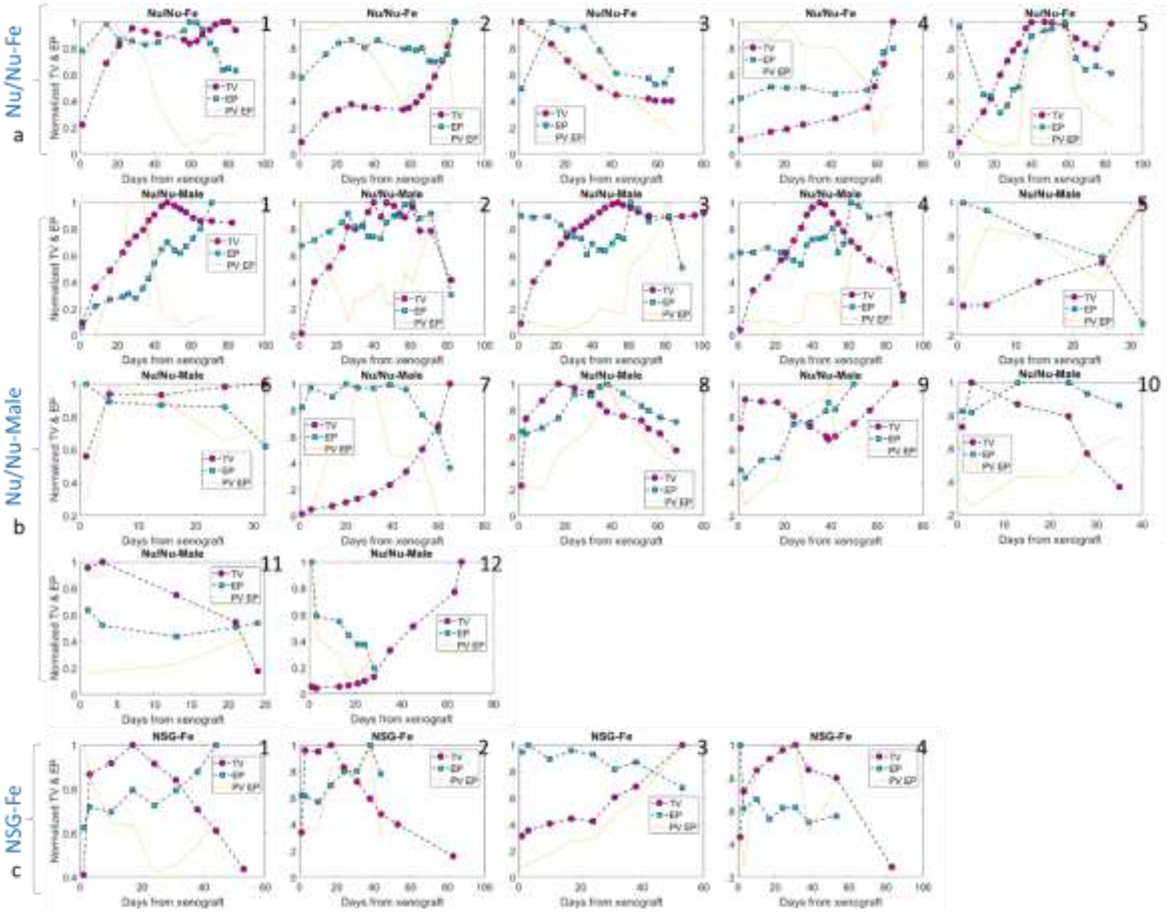


**Figure S5.** Total Volume (TV) and Average Particle Density (EP or VH. Cells) vs. Days from xenograft plot. Fig. S5(a) in first row, Fig. S4(b) in center rows and Fig. S5(c) in last rows contains data plots of Nu/Nu Female, Nu/Nu Male and NSG Female mice cohort, respectively. None of the mouse in this figure is treated. Most of the graphs show almost a linear relation being followed between both of the parameters with few exceptions(2,10,12, 15, 18). These exceptions follow inverse linear relationship.

#### 8.6 EP or VH. Cells vs. Days with treatment

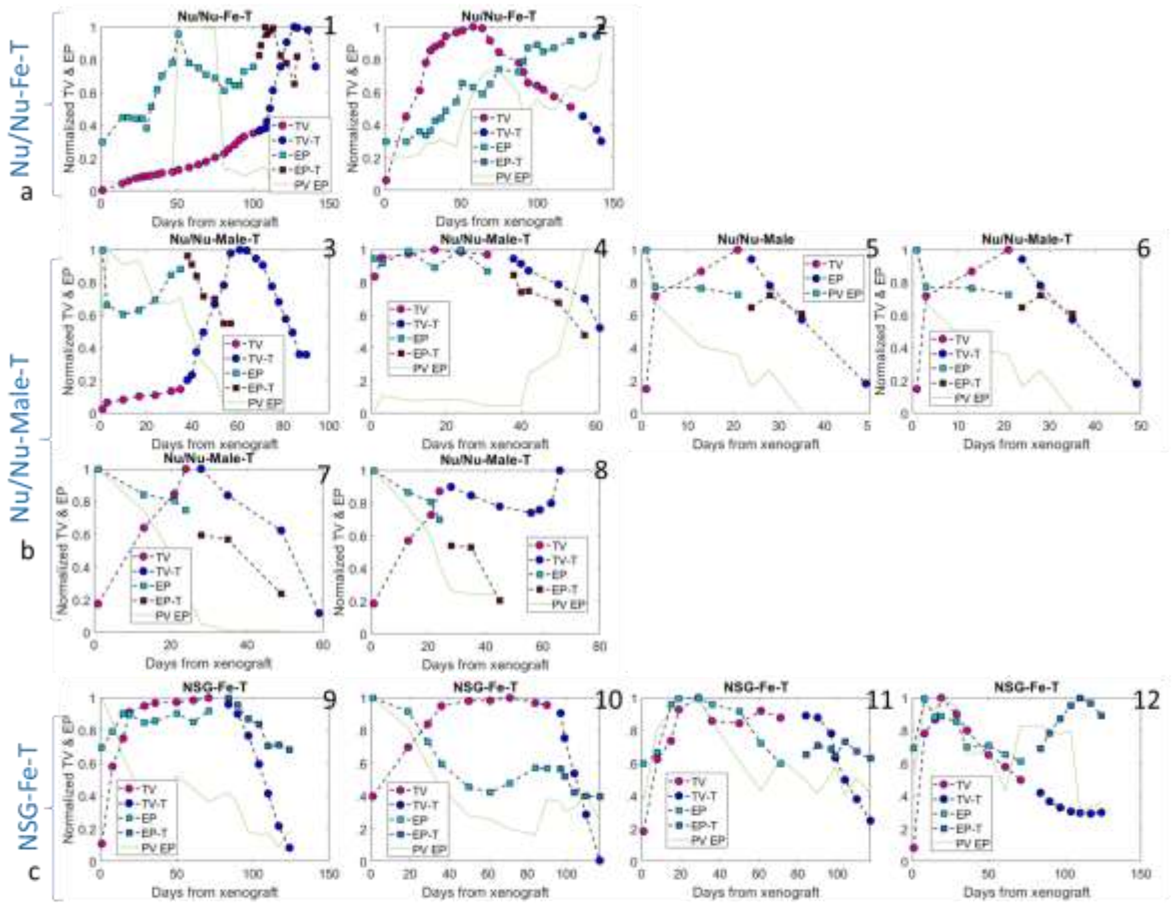


**Figure S6.** Total Volume (TV) and Average Particle Density (EP or VH. Cells) vs. Days from xenograft plot. Fig. S6(a) in first row, Fig. S6(b) in center rows and Fig. S6(c) in last rows contains data plots of Nu/Nu Female, Nu/Nu Male and NSG Female mice cohort, respectively. All of the mice in this figure have gone under nanodox dynamic photodynamic imaging assisted treatment protocol. The color scheme for marker changes once the treatment data is plotted. Before treatment again almost a linear relation seems to exists between both of the parameters with few exceptions(8,10). After treatment  inverse linear relatin is shown in few cases (1, and 12).

#### 8.7 Cases with single degree Cross correlation fit

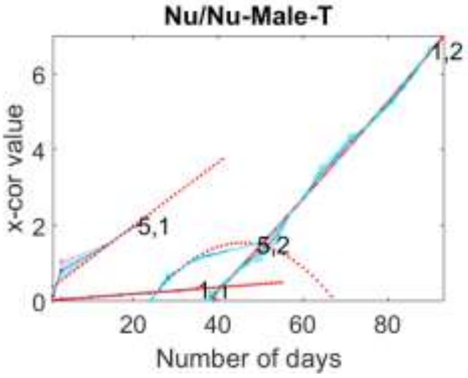


**Figure S7.** Cross correlation values between number of vitreous humor cells and tumor volume growth fit to polynomial of single degree (linear fit). The red markers are fitted values, dots in cyne color are original values and blue shade represent the error in fitting. The data belongs to Nu/Nu male mouse those have undergone the treatment protocol.

#### 8.8 Histopathology showing cells

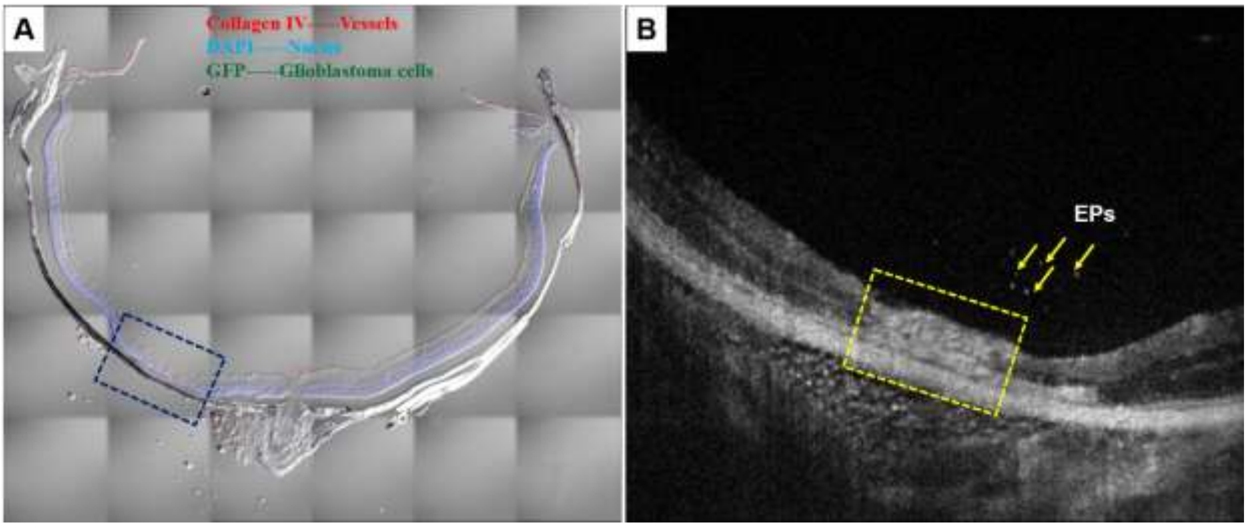


**Figure S8.** Qualitative analysis of invasive and non-invasive imaging. Invasive confocal microscopic analysis for studying immunohistochemistry. **A.** Athymic nude-foxnlnu mice (right eye), concentration - 250 cells/0.25 µl, with Nanoparticles Treatment (Nanodoxorubicin) and sacrificed on 05/11/2016. After treatment the tumor disappeared with retina atropy (region marked in dark blue). and Non-invasive OCT imaging of **B**. 2D B-scan image of Athymic nude –foxnlnu mice showing the visualization of cells in arrow and the condition of retinal atrophy in XY plane.

Table ST1: Cells growth w.r.t. age/time

<table>
<tr><th colspan="3">Cells</th><th>Function (if present)</th><th>Nu/Nu Female</th><th>Nu/Nu Male</th><th>NSGS Female(71)</th></tr>
<tr><td rowspan="5">Lymphocytes</td><td rowspan="2">T</td><td>Regulatory</td><td>Restrain NK cells</td><td rowspan="2">Increases significantly</td><td rowspan="2">Increases</td><td rowspan="2">Remains constant</td></tr>
<tr><td></td><td></td></tr>
<tr><td rowspan="2">NK</td><td>Mature/CD56$^{dim}$</td><td>Cytotoxic & anti-viral</td><td>Increases</td><td>Increases</td><td>Absent</td></tr>
<tr><td>Immature/CD56$^{bright}$</td><td>Cytokine T helper cell, regulation</td><td></td><td></td><td></td></tr>
<tr><td colspan="2">B</td><td></td><td></td><td></td><td>Absent</td></tr>
<tr><td colspan="3">Neutrophils</td><td></td><td></td><td></td><td>Absent</td></tr>
<tr><td colspan="3">Leukocytes</td><td></td><td></td><td></td><td>Absent</td></tr>
<tr><td colspan="3">Monocytes</td><td></td><td></td><td></td><td>Absent</td></tr>
<tr><td colspan="3">Microphagous</td><td>NK regulation</td><td></td><td></td><td>Defective</td></tr>
<tr><td colspan="3">T suppressor</td><td></td><td></td><td></td><td></td></tr>
<tr><td colspan="2" rowspan="2">Myeloid cells</td><td>Expressor</td><td>NK promoters</td><td rowspan="2"></td><td rowspan="2"></td><td rowspan="2"></td></tr>
<tr><td>Suppressor cells</td><td>NK inhibitors</td></tr>
</table>
